# Depletion of skin bacteria by topical antibiotic treatment accelerates onset of Zika virus disease in mice

**DOI:** 10.1101/2024.07.12.602934

**Authors:** Lana Langendries, Sam Verwimp, Sofie Jacobs, Yeranddy Aguiar Alpizar, Bert Malengier-Devlies, Rana Abdelnabi, Jeroen Raes, Chris Callewaert, Lidia Yshii, Leen Delang

## Abstract

Mosquito-borne flaviviruses have emerged as global health threats due to their rapid spread and high disease burden. The initial site of virus replication is the mosquito biting site in the host skin, yet the role of host skin bacteria remains unclear. Here, we observed accelerated progression of Zika virus (ZIKV)-induced disease in mice with depleted skin bacteria by topical antibiotic treatment prior to ZIKV infection. Increased viral loads in the blood, draining lymph node, spleen and brain of antibiotic-treated mice suggested faster ZIKV dissemination. Flow cytometry revealed increased frequencies of T cells in the lymph nodes, along with enhanced activation of T cells in the brain upon antibiotic treatment. The antibiotic-induced phenotype was rescued by restoring the skin bacteria prior to infection. Our findings demonstrate an increased susceptibility to ZIKV infection in mice with depleted skin microbiomes, suggesting a potential role of skin bacteria in modulating the infection outcome.

## Introduction

Mosquito-borne flaviviruses, such as the Zika virus (ZIKV), have emerged as significant public health concerns in recent decades, infecting approximately 400 million people worldwide annually.^1^ While ZIKV infections were previously limited to sporadic cases in Africa and South-East Asia, a major outbreak occurred on Yap Island in 2007 infecting 73% of its residents^2^, followed by larger outbreaks across the Pacific Islands in 2013. Two years later, autochthonous ZIKV infections were reported in Brazil leading to the rapid spread of ZIKV throughout the entire American continent, resulting in over 800,000 cases by January 2018.^34^

The majority of ZIKV-infected humans are asymptomatic or present mild, self-limiting febrile symptoms.^5^ Nevertheless, in most recent outbreaks ZIKV infections were increasingly associated with more severe neurological complications.^6^ A subset of infected adults develops Guillain-Barré syndrome, in which damage to the peripheral nerves leads to progressive paralysis and death in approximately 5% of cases.^7^ Additionally, vertical transmission of ZIKV can result in fetal abnormalities, such as microcephaly, designated as congenital Zika virus syndrome.^6^ In the absence of efficient mosquito control methods, vaccines and antiviral therapies, continued efforts to find new prevention and treatment strategies remain a global health priority.

Flaviviruses are inoculated by their mosquito vectors into the mammalian host skin, establishing the skin as a pivotal site for a complex triadic interplay among virus, vector and host. The presence of skin-resident and -migratory immune cells (neutrophils, monocytes, dendritic cells) allows the host to initiate a strong immune response, yet these cells can also be exploited by the virus to amplify viral replication and dissemination.^8^ Additionally, the human skin is abundantly colonized by approximately 10^6^-10^9^ bacteria per cm^2^ of skin^9,10^, exhibiting a diverse composition that varies significantly between and within individuals.^10^ While interactions between the virus and the mosquito microbiome have been studied more extensively, limited information is available regarding the influence of the host microbiome. Previous research demonstrated that the composition of the host skin microbiome influenced mosquito attraction and flavivirus transmission.^11,12^ Our previous work has shown that cell wall components of certain host bacteria, including some human skin bacteria, were able to reduce flavivirus infectivity *in vitro.*^13^ Yet, it remains uncertain whether the host skin microbiome could affect flavivirus replication in the skin and, consequently, have an impact on systemic virus dissemination and pathogenesis.

To study mosquito-borne virus pathogenesis and virus-host interactions, mouse models are frequently being used. AG129 mice (interferon (IFN) α/β/γ R-^/-^) are the standard model for ZIKV since this virus cannot efficiently antagonize the IFN response in wild type mice.^14–17^ The mouse skin microbiome can be modulated by topical application of broad-spectrum antibiotics.^18^ The effect of antibiotics on flavivirus infections has been studied before, but only by targeting intestinal or vaginal bacteria instead of skin bacteria. Flavivirus-infected mice with depleted intestinal microbiomes after oral antibiotic treatment (containing vancomycin, neomycin, ampicillin and metronidazole (VNAM)) showed higher mortality and increased viral replication. Antibiotic-treated mice also exhibited an impaired antiviral T cell response, likely contributing to the more severe infection outcome.^19^ Contrastingly, another study showed that antibiotic treatment of the vaginal mucosa with the aminoglycoside antibiotic neomycin exhibited a protective effect against ZIKV replication by increasing the expression of antiviral IFN-stimulated genes.^20^

Given that ZIKV and other flaviviruses are initially inoculated into the skin by their mosquito vectors, it is crucial to understand the impact of bacteria present at the skin inoculation site. Herein, we specifically aimed to locally reduce bacterial loads at the skin inoculation site of AG129 mice by topical application of a broad-spectrum antibiotic cream, followed by ZIKV infection. We showed that antibiotic treatment considerably accelerated progression of ZIKV-induced disease, which improved after restoring bacterial loads in the skin. Moreover, increased virus loads in the blood, draining lymph node, spleen and brain of antibiotic-treated mice suggested faster dissemination of ZIKV compared with control mice. Flow cytometry analysis revealed that antibiotic-treated mice exhibited larger frequencies of T cells in the lymph nodes, as well as enhanced activation of T cells in the brain, potentially contributed to the faster onset of neurological symptoms. Overall, our results provide evidence for a role of skin bacteria at the inoculation site in modulating the outcome of subsequent ZIKV infection in mice.

## Results

### Antibiotic treatment of the skin enhanced ZIKV disease progression in mice

To study the potential impact of host skin bacteria on ZIKV infections *in vivo*, the skin bacteriome of AG129 mice was modulated using a broad-spectrum antibiotic cream containing vancomycin, neomycin, ampicillin and metronidazole antibiotics, further referred to as VNAM cream. Since it has been described that certain antibiotics can exert an antiviral effect in addition to their anti-bacterial effect^21,22^, we first confirmed that none of the antibiotics (single or in combination) had an antiviral effect against ZIKV *in vitro* using Vero E6 cells (Fig. S1B). In the *in vivo* study, the left footpad skin of AG129 mice was treated with VNAM cream for 7 consecutive days. As oral VNAM administration was shown to affect flavivirus infectivity *in vivo*^19^, mice treated with an oral cocktail of VNAM were included as a positive control. Mice receiving vehicle cream were included as a negative control group. Treatments were well tolerated by all mice, as evidenced by the absence of weight loss and toxicity signs (Fig. S2A). After treatment, mice were subcutaneously infected in the treated footpad with ZIKV (1000 PFU) and sacrificed when showing severe weight loss or other signs of severe ZIKV-induced disease (Fig. 1B). Both oral and topical antibiotic treatment pronouncedly accelerated the ZIKV disease course, reducing the median survival from 18 days pi for vehicle-treated mice to 13 days pi for both orally and topically treated mice (Fig. 1A). Consistently, similar effects on survival were observed when mice were infected with a 10-fold lower virus inoculum (100 PFU) (Fig. S2B).

**Fig. 1.**
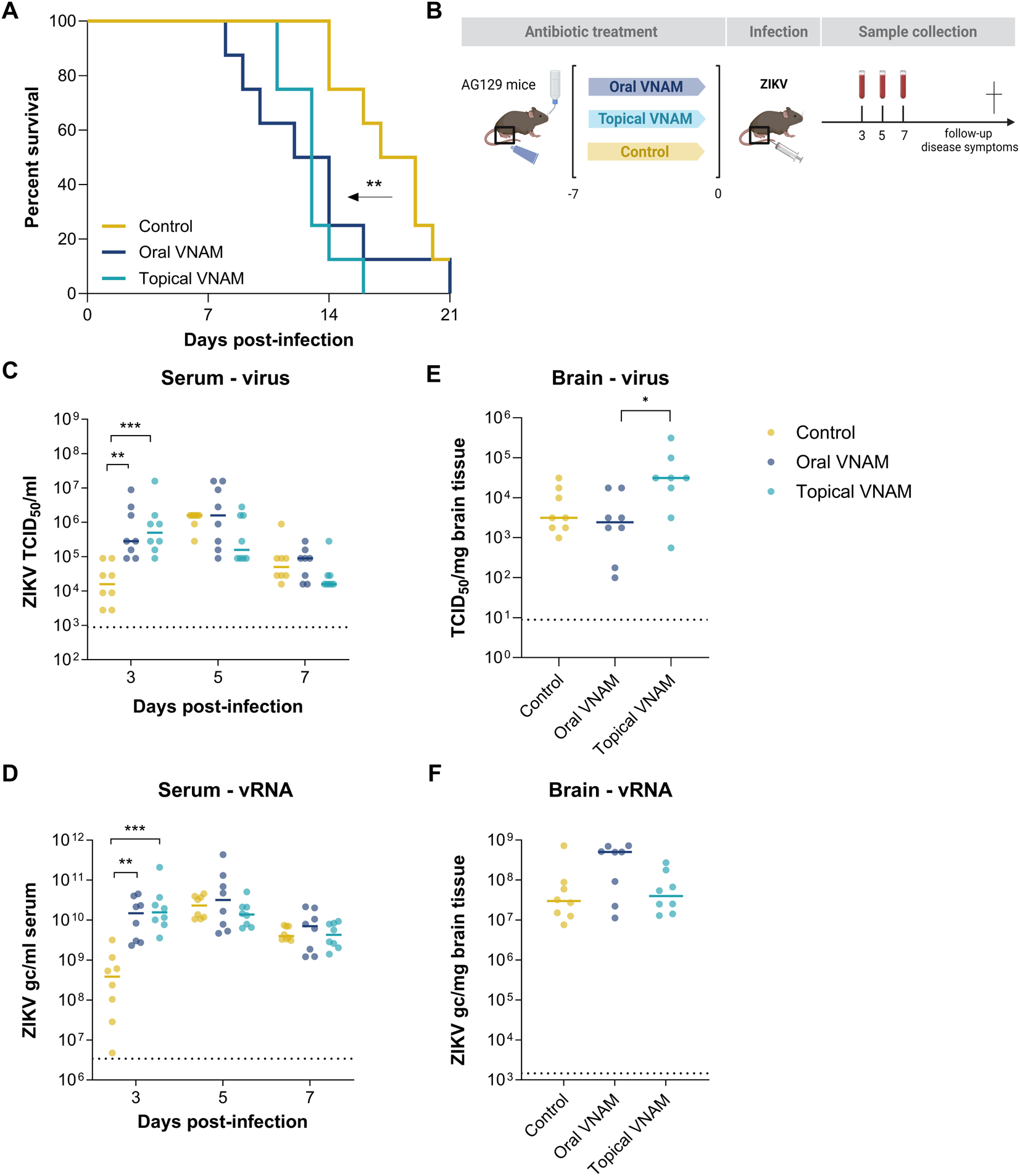
Antibiotic treatment of the skin accelerated ZIKV disease progression and increased ZIKV burden. AG129 mice (n=8 per group) were treated with an oral VNAM cocktail, an antibiotic VNAM cream, or a vehicle cream at the skin inoculation site. Subsequently, mice were subcutaneously infected in the footpad with the ZIKV PRVABC59 strain (1000 PFU). Mice were sacrificed and dissected when showing signs of severe disease. (A) Kaplan-Meier survival curves after ZIKV infection. Median values of survival were calculated and a log rank (Mantel-Cox) test was performed to assess statistically significant differences between survival curves (**, p<0.01). (B) Schematic representation of the study design. (C-D) Serum was collected on day 3, 5, 7 pi and (E-F) brain was harvested at the time of sacrifice. (C,E) Infectious virus levels were determined by end-point titrations on Vero cells and (D,F) viral RNA was quantified by qRT-PCR. (C-D) Statistical significance was assessed with a two-way repeated measures ANOVA with Sidak’s correction for multiple comparisons (**, p<0.01; ***, p<0.001). (E-F) Statistical significance was assessed with the Kruskal-Wallis with Dunn’s multiple comparisons test (*, p<0.05). (C-F) Individual data points are shown, with solid lines representing the median values. Dotted lines represent the (C,E) LOQ and (D,F) LOD. VNAM: vancomycin, neomycin, ampicillin, metronidazole; TCID50: tissue culture infectious dose 50%; vRNA: viral RNA; gc: genome copies; LOQ: limit of quantification; LOD: limit of detection.

Infectious virus and viral RNA were determined in serum collected at day 3, 5 and 7 pi. At day 3 pi, the serum of antibiotic-treated mice contained higher titers of infectious virus and viral RNA than the serum of the control group, which only reached peak viremia at day 5 pi (Fig. 1C-D). Infectious virus and viral RNA were also determined in brain tissues that were collected individually at the time each mouse had to be euthanized due to the development of severe ZIKV disease signs. A significant increase was observed in infectious virus loads in the brain of topically treated mice compared to orally treated mice (Fig. 1E-F).

### Topical antibiotic treatment resulted in local depletion of skin bacteria

To confirm whether the antibiotics were effectively inhibiting bacterial growth in the skin, skin tissue was collected after 7 days of treatment and tissue homogenates were plated on blood agar to quantify bacterial loads. VNAM treatment significantly reduced skin CFU (average reduction of 1.7 log10 CFU/skin sample) compared with the control treatment at the time of ZIKV inoculation (Fig. 2A). The principal coordinates analysis (PCoA) based on Canberra dissimilarities revealed distinct clustering of bacterial communities in the skin of VNAM-treated mice compared with control mice (Fig. 2B). Despite overlap of some samples, the overall separation between the two clusters indicated a discernible shift in microbiome composition upon VNAM treatment. Similar observations were made using Jaccard and Bray-Curtis metrics (Fig. S3 A, B). To investigate a potential systemic effect of the topical antibiotics, we quantified 16S genome copies of bacteria in feces before and after treatment. As expected, the oral antibiotic treatment caused a bacterial reduction in the feces, resulting in a 1.6 log_10_ decrease in 16S genome copies/mg feces (Fig. 2C). In contrast, topical VNAM treatment did not affect 16S genome copies in the feces. To fully exclude a potential systemic effect of the topical antibiotic treatment, bacterial diversity was investigated by 16S rRNA sequencing on the feces samples. Bacterial community composition was not significantly affected by the topical antibiotic treatment as measured by the alpha-diversity Shannon index (Fig. 2D).

**Fig. 2.**
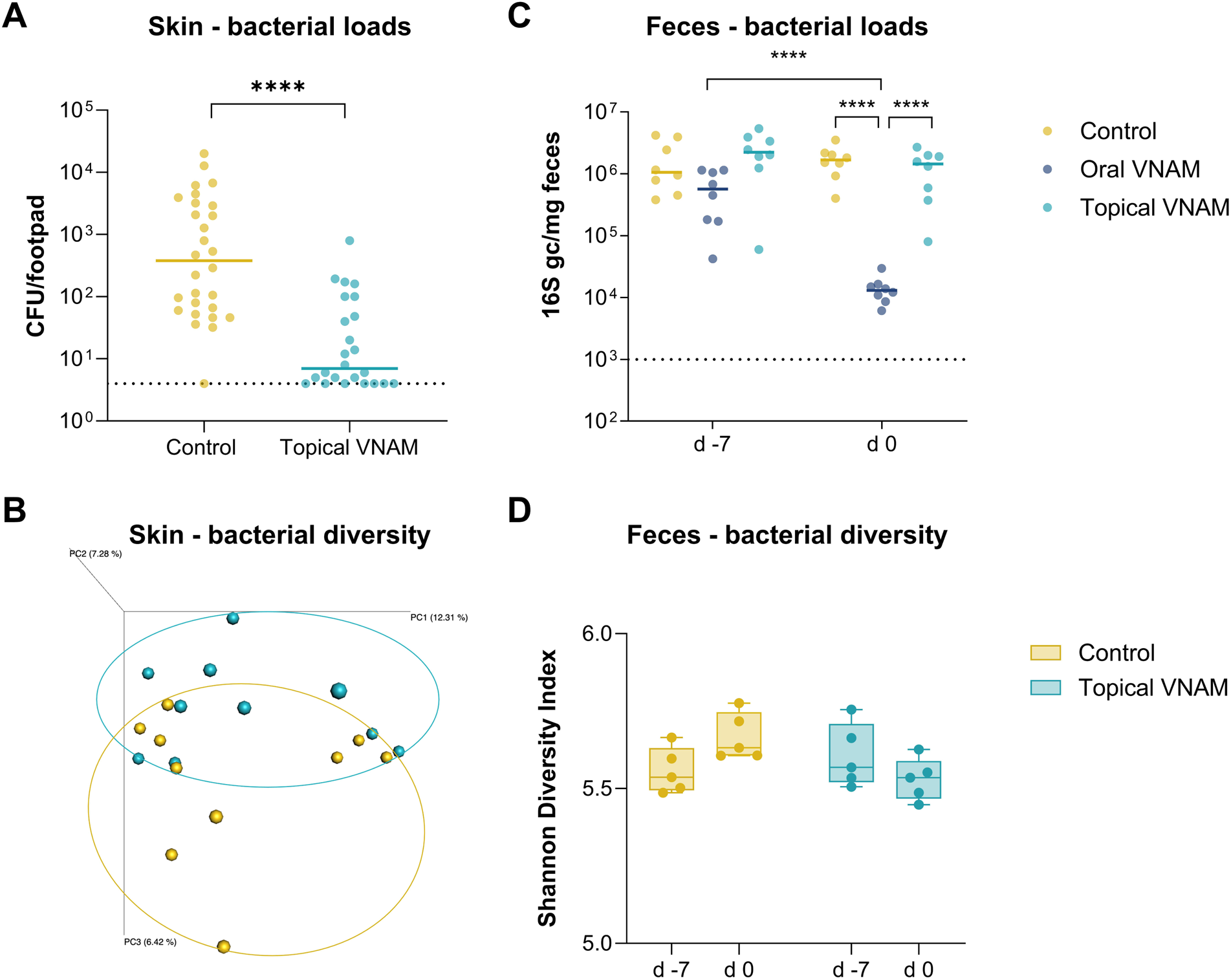
Topical antibiotic treatment locally depleted bacteria in the skin. Mice were orally or topically treated with VNAM or vehicle. (A) Skin samples or (B) skin swabs were collected after 7 days of treatment and (C, D) feces samples were collected both before and after 7 days of treatment. (A) CFUs were determined after plating the skin homogenates on blood agar. Data are representative of three independent experiments (Control: n=28, VNAM: n=24). (B) Beta-diversity based PCoA plots (Canberra metrics) of skin bacterial communities after 16S metagenomic sequencing of skin swabs (n=10 per group) X, Y, Z axes show PC1, PC2 and PC3 components which explain 12.31 %, 7.28 % and 6.42 % of variation, respectively. (C) 16S genome copies were quantified in feces samples by 16S qRT-PCR (n=8 per group). (D) Shannon diversity index representing alpha-diversity of bacterial communities in feces after 16S metagenomic sequencing (n=5 per group). (A) Statistical significance was assessed with a Mann-Whitney U test (****, p<0.0001). (C, D) Statistical significance was assessed with a two-way repeated measures ANOVA with Sidak’s correction for multiple comparisons (ns, p>0.05; ****, p<0.0001). (A,C) Individual data points are shown, with solid lines representing the median values. The dotted lines represent the LOD. (D) Data is represented as floating bars, showing the individual data points with solid lines indicating the median values. CFU: colony forming units; gc: genome copies.

### Topical antibiotic treatment affected the viral burden in ZIKV target tissues

Given the increased susceptibility of VNAM-treated mice to ZIKV infection, we next aimed to investigate whether topical VNAM treatment could affect initial viral replication in the skin and/or dissemination to secondary tissues. Therefore, we first characterized the replication kinetics of ZIKV in different target tissues upon inoculation into the footpad of untreated AG129 mice. Viral RNA levels started to increase in the skin from 8 h pi onwards, reaching up to 6.4 log_10_ genome copies/mg at day 3 pi, showing active viral replication (Fig. S4A). Additionally, ZIKV was able to spread to the draining lymph node, serum and spleen, as demonstrated by increasing viral RNA levels within the first 3 days pi (Fig. S4B-D).

Next, we studied the replication kinetics in mice that were topically treated with VNAM or control cream prior to ZIKV infection. Based on the previous study (Fig. S4), tissues were analyzed at 16 h and 48 h pi. The viral RNA load was significantly elevated (∼0.8 log_10_ increase) in the inguinal lymph nodes of mice treated with VNAM cream compared with mice treated with control cream at 48 h pi (Fig. 3D). No significant differences in viral RNA in the skin, serum or spleen were detected between VNAM- and vehicle-treated groups. Infectious virus levels were not significantly different either at these early time points (Fig. 3). The control cream itself was not influencing viral replication, since no differences in viral loads were observed between untreated mice and control cream-treated mice (Fig. S5A-G).

**Fig. 3.**
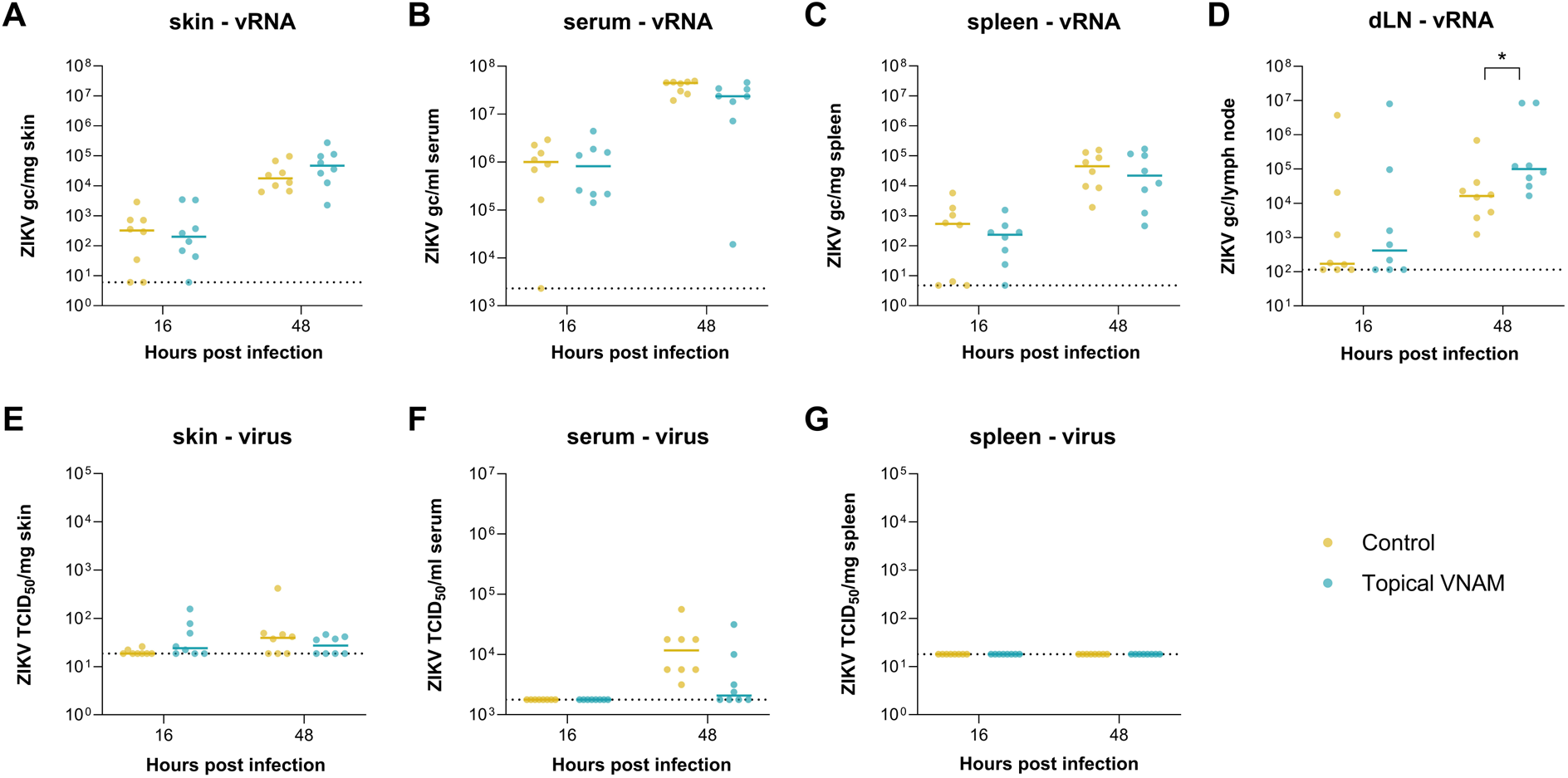
Topical antibiotic treatment increased early ZIKV burden in the draining lymph node. Mice were topically treated with VNAM or vehicle (n=8 per group) prior to infection with ZIKV (1000 PFU). Tissues were analyzed at 16 and 48 h pi. ZIKV RNA levels in (A) skin of the inoculation site, (B) serum, (C) spleen, (D) draining lymph node, as determined by qRT-PCR. TCID_50_ values in (E) skin of the inoculation site, (F) serum and (G) spleen, quantified by viral end-point titrations on Vero cells. Individual data points are shown, with solid lines representing the median value. Statistically significant differences between control or VNAM-treated mice were assessed with a two-way ANOVA with Sidak’s correction for multiple comparisons (*, p<0.05). (A-D) Dotted lines represent the LOD. (E-G) Dotted lines represent the LOQ. vRNA: viral RNA; gc: genome copies; pi: post infection; TCID50: tissue culture infectious dose 50; LOD: limit of detection; LOQ: limit of quantification.

Viral loads were also measured in target tissues at later time points in the infection course to assess the effect on systemic pathogenesis (Fig 4). At day 7 pi, a significant elevation of infectious virus titers (∼0.7 log_10_ TCID50/mg), but not of viral RNA, was observed in the spleen of VNAM-treated mice (Fig. 4A, D). Although not statistically significant, we also observed an increasing trend in the infectious virus levels in the brain and spinal cord (∼1 log_10_ TCID50/mg) at day 9 pi following VNAM treatment (Fig. 4E-F).

**Fig. 4.**
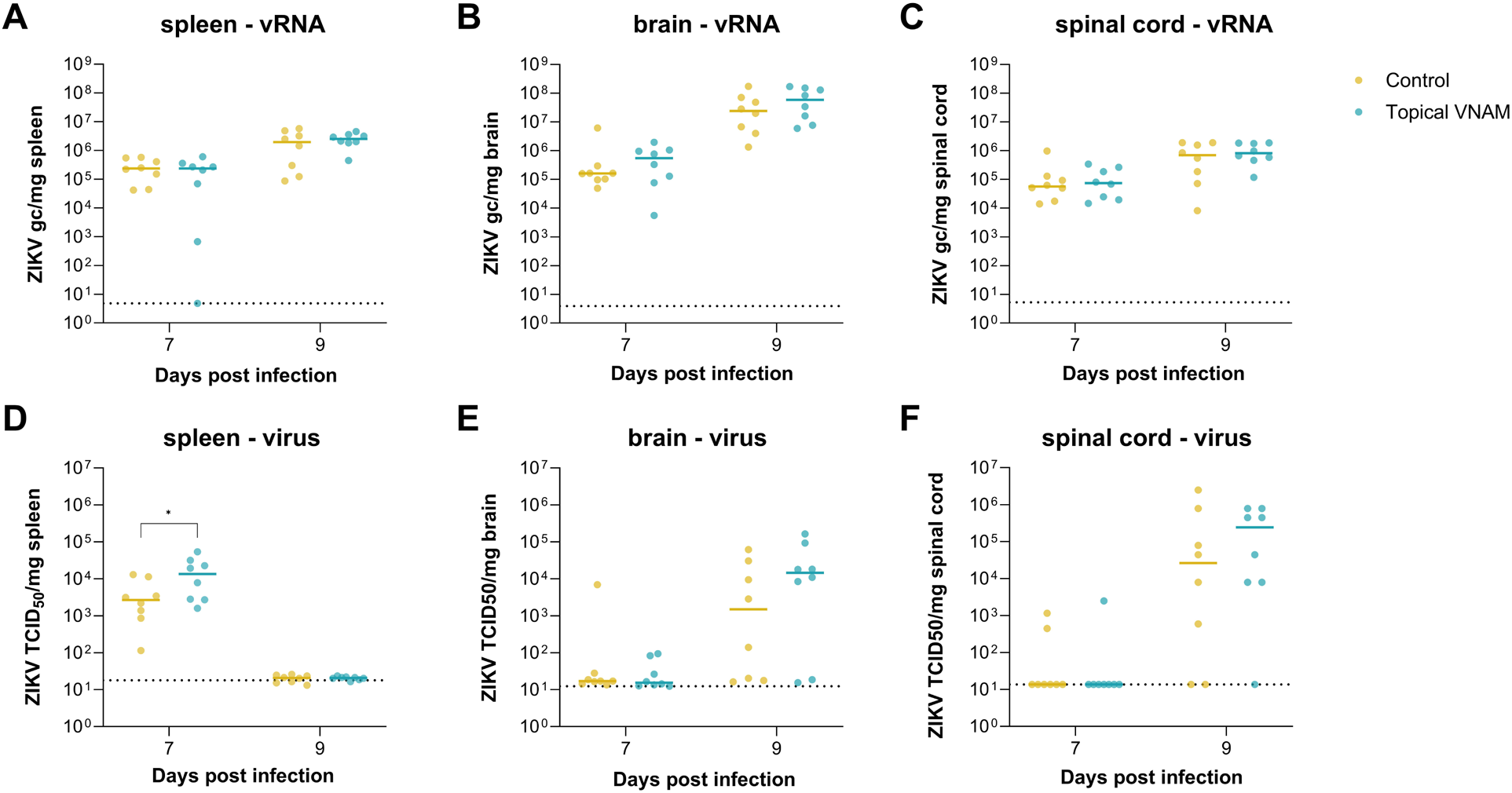
Antibiotic treatment of the skin increased late ZIKV titers in the spleen. Mice were topically treated with VNAM or vehicle cream (n=8 per group) prior to infection with ZIKV (1000 PFU). Tissues were analyzed at day 7 and 9 pi. ZIKV RNA levels in (A) spleen, (B) brain, (C) spinal cord, determined by qRT-PCR. TCID_50_ values in (D) spleen, (E) brain and (F) spinal cord, quantified by viral end-point titrations on Vero cells. Individual data points are shown, with solid lines representing the median values. Statistical significance was assessed with a two-way ANOVA with Sidak’s correction for multiple comparisons (*, p<0.05). (A-C) Dotted lines represent the LOD; (D-F) dotted lines represent the LOQ. vRNA: viral RNA; gc: genome copies; pi: post infection; TCID50: tissue culture infectious dose 50; LOD: limit of detection; LOQ: limit of quantification.

### Immune cell composition in the skin and draining lymph node were altered in antibiotic-treated mice

Since it is known that antibiotics can exert immunomodulating effects^19,20,23^, flow cytometry was performed on the footpad skin of uninfected or ZIKV-infected mice treated priorly with control or VNAM cream to determine the immune cell composition. Skin samples were collected at both 16 h and 48 h pi. Different immune cell types residing in or migrating to the skin were evaluated based on the expression of CD45, CD3, CD19, CD11b, CD11c, CD64, Ly6C, MHC-II and CD207. The gating strategy used is illustrated in Fig. S6. The frequencies of the different cell types (Langerhans cells, conventional type 1 dermal DCs (cDC1), monocyte-derived DCs, monocytes, CD11b+ classical DCs, dermal macrophages, granulocytes, T and B cells) were determined relative to the total number of CD45+ cells (Fig. 5A-I). During the infection course, percentages of certain immune cells such as CD11b+ cDCs and monocyte-derived DCs significantly elevated, indicating that these immune cells were recruited to the skin upon ZIKV infection. Contrastingly, reduced percentages of immune cells such as LCs and dermal DCs point towards migration and possible dissemination of these cells out of the skin. Importantly, the frequency of monocytes decreased in the skin of VNAM-treated mice and was significantly lower than those of vehicle-treated mice at 48 h pi (Fig. 5D).

**Fig. 5.**
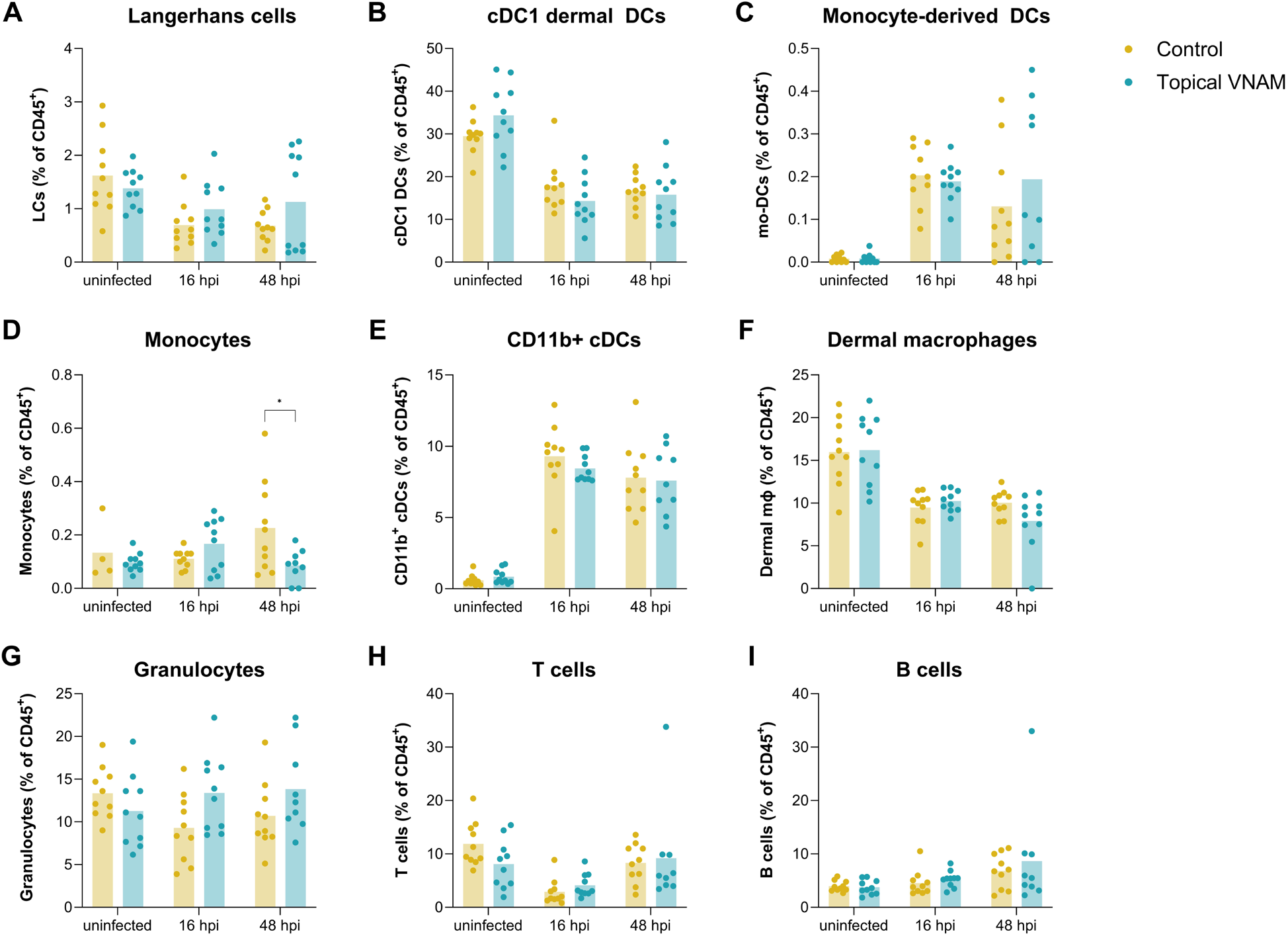
Immune cell composition in the skin was altered following topical antibiotic treatment. Mice were topically treated with VNAM or vehicle (n=10 per group). Footpad skin tissue was collected from uninfected mice or mice infected with ZIKV (1000 PFU) at 16 and 48 h pi to perform flow cytometry. Bar plots showing the mean and individual data points of the percentages of (A) Langerhans cells, (B) cDC1 dermal DCs, (C) monocyte-derived DCs, (D) monocytes, (E) CD11b+ cDCs, (F) dermal macrophages, (G) granulocytes, (H) T cells and (I) B cells among CD45^+^ cells. Statistical significance was assessed with a two-way ANOVA with Sidak’s correction for multiple comparisons (*, p<0.05). DC: dendritic cells; hpi: hours post infection.

As the viral burden was significantly increased in the lymph nodes of VNAM-treated mice at 48 h pi (Fig. 4B), immune cell composition was also assessed in the lymph nodes by flow cytometry. Different cell types were evaluated based on the expression of CD45, TCRβ, CD4, CD8, CD19, NK1.1, CD11c, Ly6C, Ly6G and SiglecF. The gating strategy is illustrated in Fig. S7. The frequencies of different cell types (CD4+ and CD8+ T cells, B cells, DCs, monocytes, neutrophils and natural killer cells) were determined relative to the total number of CD45+ cells (Fig. 6A-G). Topical treatment with VNAM resulted in a more rapid increase of CD4+ T and CD8+ T cells early in the infection course, as their percentage was significantly higher compared to that in vehicle-treated mice at 16 h pi (Fig. 6A-B). Later in the infection, VNAM-treated mice exhibited significantly lower frequencies of neutrophils (at day 7 and 9 pi) and NK cells (at day 7 pi) (Fig. 6F-G).

**Fig. 6.**
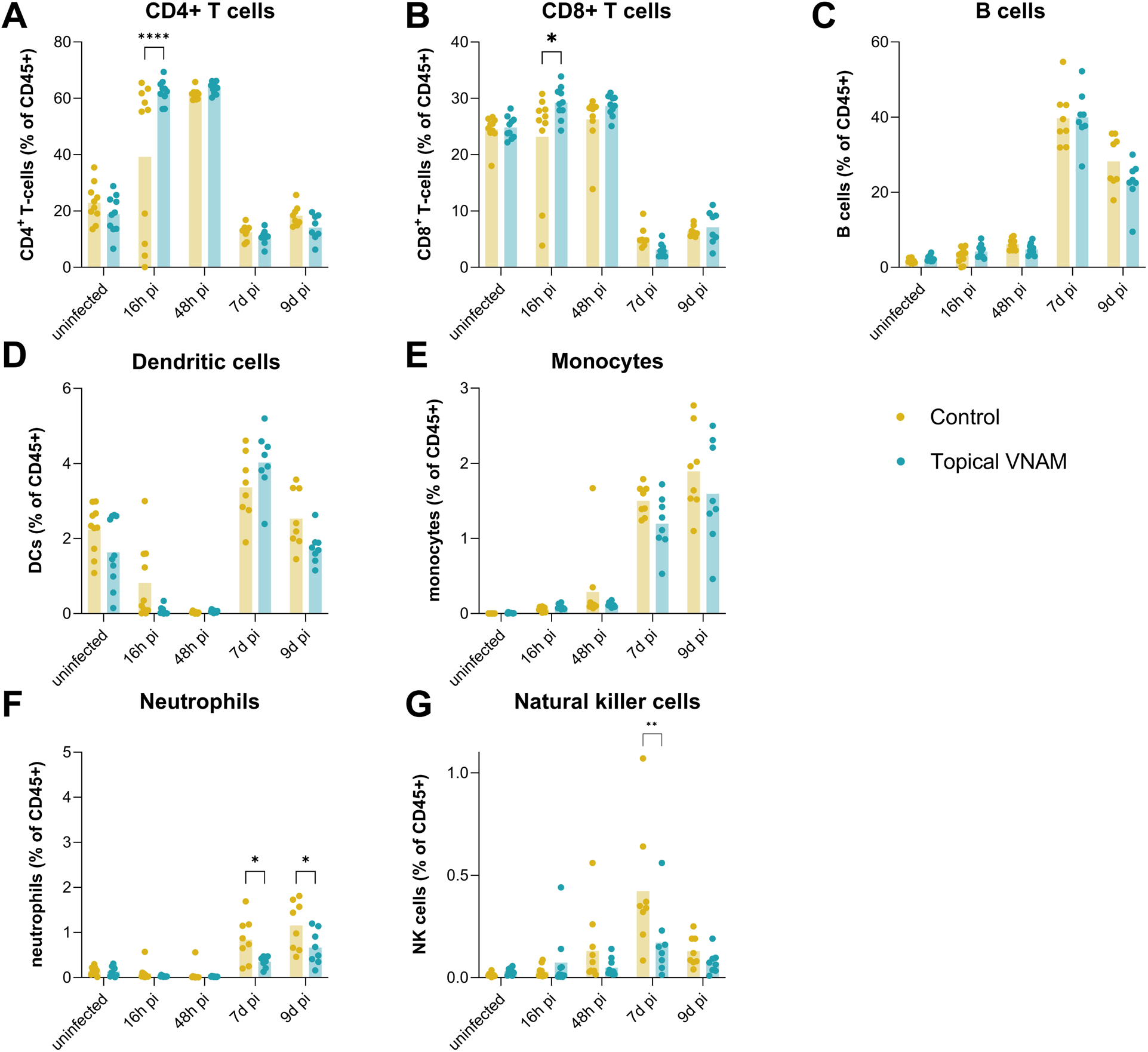
Immune cell composition in the skin was influenced by topical antibiotic treatment. Mice were topically treated with VNAM or vehicle (Uninfected, 16 h pi, 48 h pi: n=10 per group; 7d pi, 9d pi: n=8 per group). The draining inguinal lymph node was collected from uninfected mice or mice infected with ZIKV (1000 PFU) at 16 and 48 h pi or 7 and 9 d pi to perform flow cytometry. Bar plots showing the mean and individual data points of the percentages of (A) CD4+ T cells, (B) CD8+ T cells, (C) B cells, (D) dendritic cells, (E) monocytes, (F) neutrophils and (G) natural killer cells among CD45+ cells. Statistical significance was assessed with a two-way ANOVA with Sidak’s correction for multiple comparisons (*, p<0.05; **, p<0.01; ****, p<0.0001). hpi: hours post infection.

### Antibiotic treatment of the skin increased activation of T-cells in the brain

Given the increased frequencies of T cells in the draining lymph nodes and their importance in ZIKV neuropathogenesis^24–26^, we studied the T cell population in the brain by flow cytometry. At day 7 pi, T cells were analyzed based on the expression of CD4, CD8, CD69, CD44 and CD62L. The gating strategy is illustrated in Fig. S8. Expression of the activation marker CD69 was significantly elevated in the brains of VNAM-treated mice, with increases in mean fluorescence intensitys of 328 and 582 counts, respectively, on CD4+ and CD8+ T cells (Fig. 7A-B). Additionally, a small but significant increase in effector memory CD8+ T cells, relatively to the total CD4+ and CD8+ population, was observed following VNAM treatment, while no increase was measured in memory T cells (Fig. 7C).

**Fig. 7.**
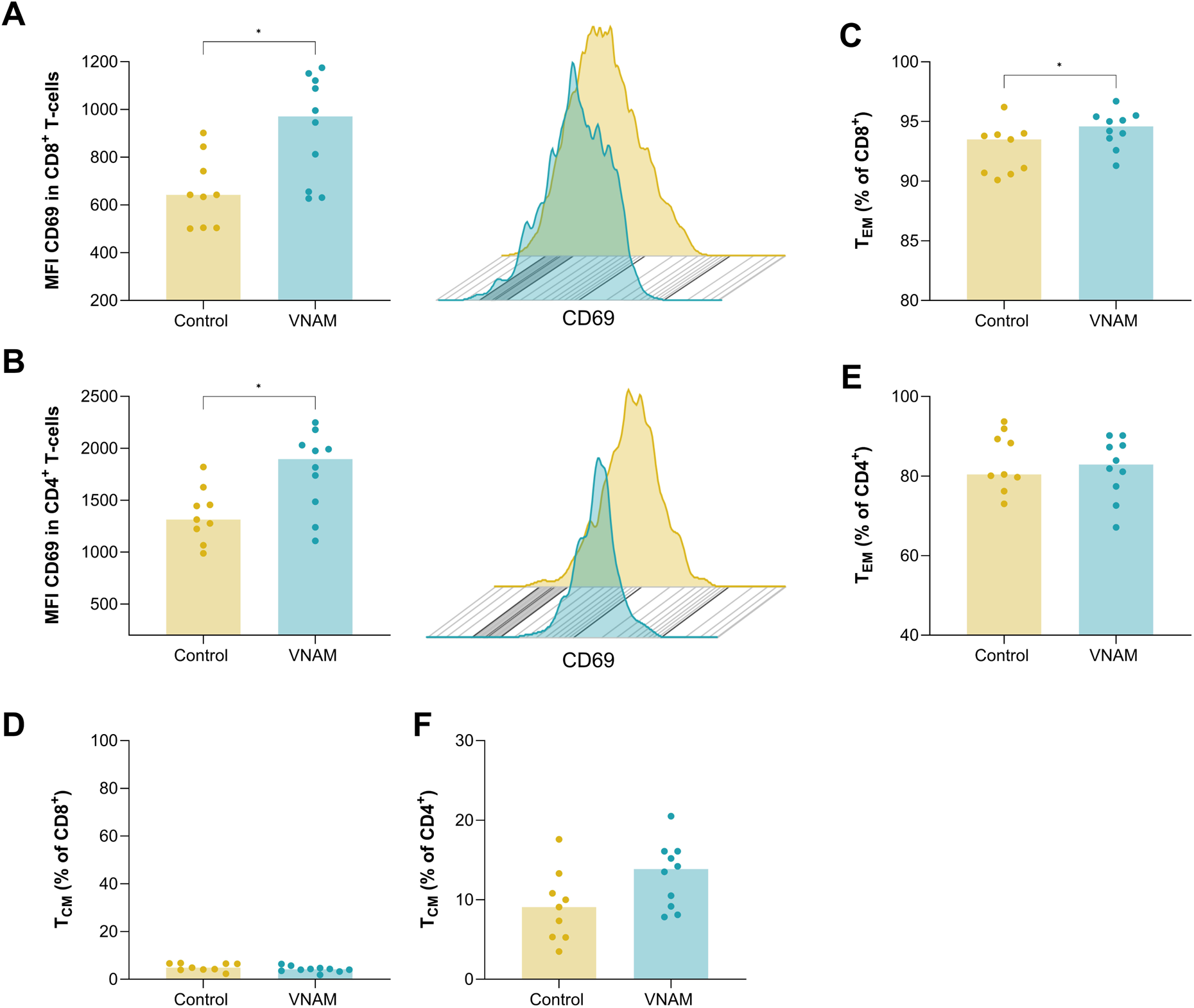
Topical antibiotic treatment increased activation of T cells in the brain. Mice were topically treated with VNAM (n=10) or vehicle (n=9) and infected with ZIKV (1000 PFU). The brain was collected at day 7 and 9 pi to perform flow cytometry. (A-B) Bar plots showing mean and individual data points of the mean fluorescence intensity of CD69 in (A) CD8+ and (B) CD4+ T cells. Representative histogram of CD69 expression in one sample per condition is shown. (C-F) Bar plots showing the mean and individual data points of percentages of (C) effector memory CD8+ T cells, (D) central memory CD4+ T cells, (E) effector memory CD4+ T cells, (F) central memory CD4+ T cells, relative to the total percentage of CD8+ and CD4+ T cells. Statistical significance was assessed with a Mann-Whitney U test (*, p<0.05). MFI: mean fluorescence intensity; T_EM_: effector memory T cells; T_CM_: central memory T cells.

### Topical antibiotic treatment influenced muscle-but not immune-related gene expression

To further unravel the mechanism underlying the enhanced progression of ZIKV-induced disease after topical VNAM treatment, we compared gene expression in the skin between VNAM- and vehicle-treated mice by bulk RNA sequencing of skin tissue. The principal component analysis (PCA) plot showed that most of the VNAM samples clustered together, whereas we could not detect clear clustering of the samples in the control group (Fig. S9A). We identified a total of 215 genes that were differentially expressed by at least a two-fold difference and an FDR-corrected p-value of <0.1^27^, from which 212 genes were downregulated and only 3 genes were upregulated upon VNAM treatment (Fig. S9B). Gene ontology and ingenuity pathway analysis revealed that 47 of these downregulated genes (e.g. Myl3, Actn2, Clcn1 genes) were involved in muscle contraction processes (Fig. S9D and Table S1). However, no immune-related genes were differentially expressed upon VNAM treatment of the skin.

### Restoring skin bacteria rescued ZIKV phenotype upon topical antibiotic treatment

Finally, we assessed whether the antibiotic-induced ZIKV phenotype could be rescued by restoring the skin bacteria after halting the antibiotic treatment. To this end, mice were treated with VNAM cream on the footpad skin for 7 days, followed by a period of 3 days during which no treatment was given, prior to the ZIKV infection (VNAM 3 day stop group) (Fig. 8A). The 3 days period without treatment allowed bacteria to recolonize the skin (Fig. 8B), even exceeding bacterial loads in the skin of control mice (increase of 2.8 log_10_ CFU/skin sample compared with skin of VNAM-treated mice and 1.1 log_10_ CFU/skin sample compared with skin of vehicle-treated mice). Following ZIKV infection, both oral and topical VNAM treatment (median survival of 10 days) significantly accelerated the progression of ZIKV-induced disease compared with control treatment (median survival of 13 days) (Fig. 8C; p = 0.003 for topical VNAM vs control; p = 0.006 for oral VNAM vs control). However, when allowing recolonization of skin bacteria, the median survival significantly increased again (median survival of 12.5 days) (p= 0.02; oral/topical VNAM vs VNAM stop), and was similar to mice in the control group (p=0.4 VNAM stop vs control). These data suggest that the increased susceptibility to ZIKV was directly due to the local reduction in skin bacteria.

**Fig. 8.**
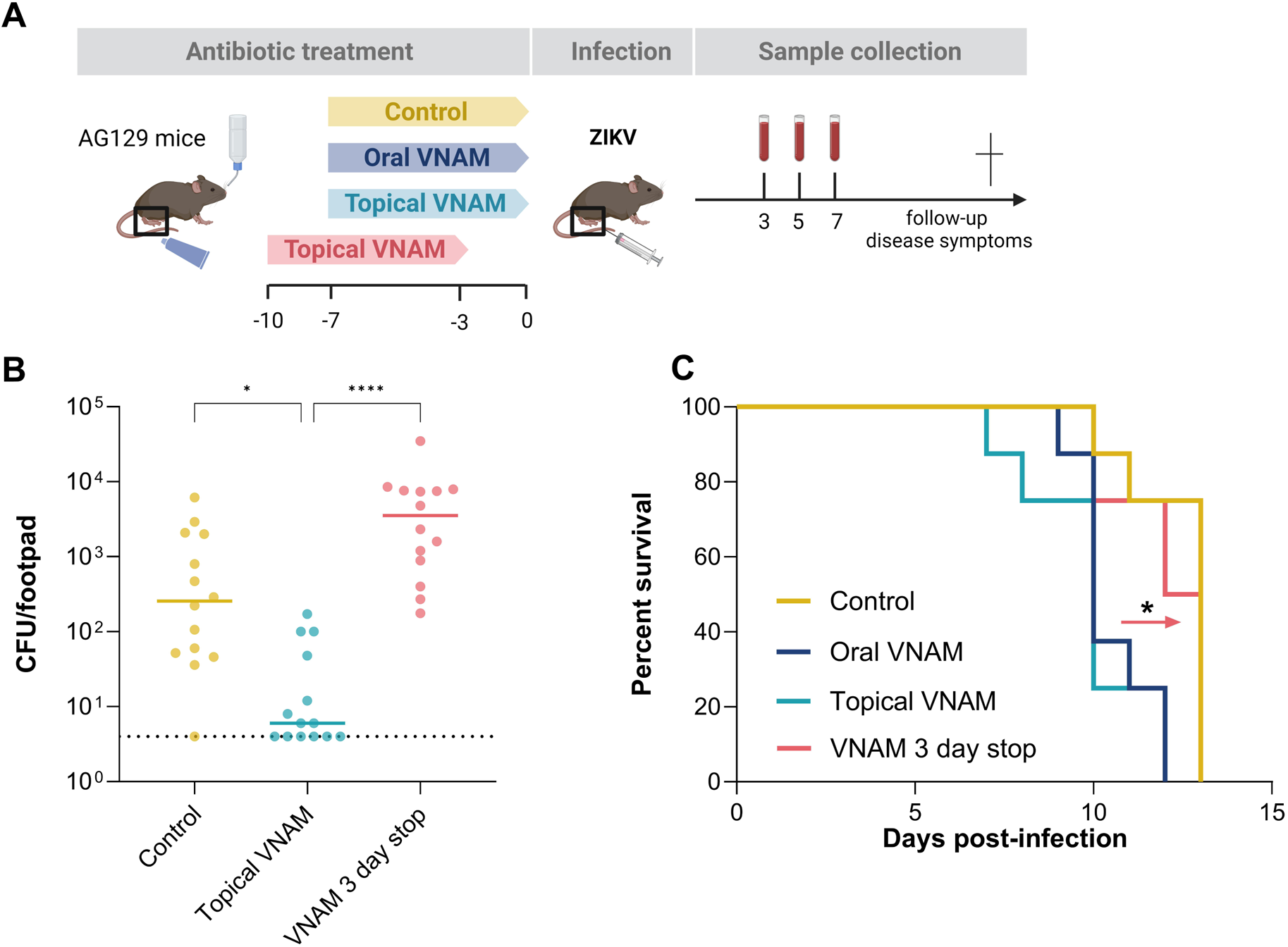
Restoring bacteria levels in the skin prior to infection rescued antibiotic-induced susceptibility to ZIKV. (A) Schematic representation of the study design. Female AG129 mice were treated with an oral VNAM cocktail, a VNAM cream or a vehicle cream at the footpad skin for 7 days prior to ZIKV infection (1000 PFU). An additional group received similar treatment with VNAM cream for 7 days which was stopped 3 days prior to the infection. Blood was collected at day 3, 5 and 7 pi. Mice were sacrificed and dissected when showing signs of severe diseases. (B) Skin samples were harvested after treatment or after the stop period and CFUs were determined after plating the skin homogenates on blood agar. Data are shown from two independent experiments (n=14 per group). Statistical significance was assessed with the Kruskal Wallis test with Dunn’s multiple comparisons test (*, p<0.05; ****, p<0.0001). (C) Kaplan-Meier survival curves after ZIKV infection (n=8 per group). Median values of survival were calculated and a log-rank (Mantel-Cox) test was performed to assess statistically significant differences between survival curves (*, p<0.05). CFU: colony forming units.

## Discussion

There is an urgent need for strategies that reduce the burden of mosquito-borne viruses. The initial inoculation of a mosquito-borne virus in the skin is an essential step to establish a productive infection in the host. Whether the presence of bacteria at the skin could affect mosquito-borne virus replication, dissemination and ultimately, pathogenesis in the host, has not been studied in detail. In this study, we locally adapted the skin bacteria of AG129 mice prior to ZIKV infection by topical treatment with a broad-spectrum antibiotic cream. Herein, we demonstrated that antibiotic-treated mice exhibited an increased susceptibility to ZIKV disease, suggesting a potential role of host skin bacteria in modulating the outcome of ZIKV infections.

Previous studies showed that the murine skin is predominantly colonized by Gram-positive *Staphylococcus*, *Streptococcus*, *Enterococcus*, *Corynebacterium* and *Bacillus,* and Gram-negative anaerobic *Alistipes* spp.^28,29^ To ensure efficient inhibition of bacterial growth, the antibiotic cream used in our studies was composed of four antibiotics including vancomycin, neomycin, ampicillin and metronidazole, selected for their known broad-spectrum activity. In general, vancomycin is active against Gram-positive cocci and bacilli, such as *Staphylococcus*, *Streptococcus* and *Corynebacterium* spp.^30^; neomycin has activity against the majority of Gram-negative and a few Gram-positive bacteria^31^; ampicillin is effective against many Gram-positive (*Staphylococcus, Streptococcus*, *Enterococcus*) and medically important Gram-negative bacteria (*Escherichia coli*, *Haemophilus influenzae*)^32,33^ and metronidazole is active against Gram-positive and Gram-negative anaerobic bacteria.^34^ This combination of antibiotics allowed us to effectively reduce local bacteria as evidenced by a marked decrease in bacterial loads in the footpad skin (Fig. 2A). In addition, this broad-spectrum antibiotic cocktail was used in a previous study as an oral treatment to reduce the intestinal microbiome.^19^ Consistent with the broad-spectrum activity of the antibiotics, 16S sequencing of the skin samples confirmed that VNAM treatment induced a shift in the composition of skin bacterial communities. Variability within antibiotic-treated skin samples likely reflected variable susceptibility to the antibiotics of different bacteria in the skin. Likewise, variability was observed within the control group, highlighting the natural heterogeneity that is associated with the skin microbiome.^9^ Experiments with larger sample sizes would be required to fully capture the complexity and variability observed in bacterial responses to VNAM treatment.

It is crucial to acknowledge that complete eradication of skin bacteria was not achievable due to the inherent exposure of the footpad skin to various environmental sources of contamination (i.e. bedding material, feces, cage mates, …). To address this limitation, future studies could consider utilizing germ-free mice to entirely eliminate influences of background bacteria on the experimental outcomes, although this would also come with deficiencies in the immune system.^35^ Nevertheless, the observed reduction in bacteria in this study was sufficient to reveal significant effects on ZIKV infection and disease outcome, highlighting the importance of considering not only the presence or complete absence of bacteria but also their relative abundance in determining host-pathogen interactions.

Topical VNAM treatment prior to ZIKV infection severely affected the disease progression in mice, reducing median survival by 5 days (Fig. 1A). This effect was consistent even when using a lower inoculum of ZIKV (Fig. S2B) and was not due to any toxic effect of the antibiotic treatment (Fig. S2A). Similar findings have been reported in a previous study where enhanced susceptibility to flavivirus infections was observed in mice whose intestinal microbiome was depleted with oral VNAM treatment^36^. To address the possibility of systemic bacterial depletion as a contributing factor in our studies, considering potential oral uptake of the antibiotic cream, we collected feces before and after treatment to assess bacterial loads and composition. Our analysis revealed no significant reduction in bacterial abundance nor in bacterial diversity in the feces of mice treated topically with VNAM cream (Fig. 2C, D). This finding provided evidence that the observed effects on ZIKV disease in our study were primarily attributed to the targeted reduction of local bacteria in the skin, rather than to a systemic alterations in bacteria levels and/or composition. The fact that similar effects were observed with both oral and topical VNAM treatment supports a broad role of bacteria across various microbial sites of the host in modulating flavivirus infections, albeit through distinct mechanisms.

We next explored potential underlying mechanisms by which alteration of the skin bacteriome could influence ZIKV infections. We showed that VNAM treatment led to increased viral loads in the draining lymph node and serum at day 2 and 3 pi, respectively, suggesting that ZIKV facilitated dissemination in treated mice compared with untreated mice (Fig. 1C, 3D). This was supported by increased infectious virus titers in the brain at the time of sacrifice, although no difference was seen in viral RNA levels (Fig. 1E-F). As the time of sacrifice and thus of brain collection varied between VNAM- and vehicle-treated mice, it is plausible that there might have been a more significant difference in viral loads if brain samples would have been collected simultaneously. We therefore assessed the effects on systemic pathogenesis and measured viral loads at later time points in the brain, spinal cord and spleen, key organs in ZIKV-induced disease. In the spleen of VNAM-treated mice, infectious virus titers were significantly elevated at day 7 pi (Fig. 4D). Similar trends were observed in the brain and spinal cord (Fig. 4E-F). Although not statistically significant, the higher virus burden could be associated with enhanced neuropathogenesis and lethality, as previously shown by ^37^.

Previous studies have highlighted the multifaceted impact of antibiotics and the corresponding depletion of bacteria on the host immune response. For instance, oral antibiotic treatment was shown to affect not only survival and viral load in flavivirus-infected mice, but also to diminish CD8+ T cell levels in the spleen and lymph node^19^. Furthermore, an oral cocktail of vancomycin and ampicillin was associated with an impaired IFN response, thereby increasing the susceptibility of circulating monocytes to alphaviruses and promoting systemic viral dissemination.^36^ In this context, our study investigated whether antibiotic treatment and depletion of host bacteria in the skin affected immune responses upon ZIKV infection. We observed a local reduction in the frequency of monocytes at the skin inoculation site of VNAM-treated mice at 48 h pi (Fig. 5D). It is conceivable that the reduced local monocyte frequency reflects a more rapid migration of these cells to systemic sites such as the lymph nodes and blood. Given that monocytes can become infected with ZIKV in the skin^38,39^, they could in this way have accelerated virus dissemination in the host. Alternatively, since monocytes are pivotal in the antiviral immune response^40,41^,their reduced presence could have facilitated viral replication, consistent with the observed higher viral loads in the blood and lymph nodes of VNAM-treated mice. This is also in line with the reduced levels of neutrophils and NK cells in the lymph nodes later in the infection course (Fig. 6F-G), both of which are key effector cells in the antiviral immune response. Previous studies have demonstrated their protective roles against ZIKV disease by promoting immune-mediated virus elimination.^42–44^ Also for DENV, the NK cell response was identified as an important factor influencing disease severity.^45^ The overall diminished antiviral immunity observed following antibiotic treatment likely contributed to the accelerated ZIKV disease progression.

Interestingly, we observed an increased frequency of CD4+ and CD8+ T cells in the draining lymph nodes upon VNAM treatment early in the infection course (Fig. 6A-B). In immunocompetent mice, ZIKV infections are characterized by a robust influx of CD8+ T cells into the central nervous system. Their antiviral activity controls local virus replication, depending on their number and differentiation status.^25,26^ The role of CD8+ T cells in IFN-deficient mice upon ZIKV infection is more controversial. Adoptive transfer of CD8+ T cells from immunocompetent mice to IFNAR mice was found to reduce virus loads in the brain and spinal cord.^46^ Consistently, depletion of CD8+ T cells in IFNAR mice increased infection of neurons and virus burden in the brain. However, these mice also showed decreased development of paralysis and improved survival, showing that these cells are not only associated with viral clearance but also with increased neuropathogenesis. Their antiviral activity aiming at virus elimination induced lysis of ZIKV-infected neurons; however, as a consequence this also mediated ZIKV-associated neuropathogenesis leading to paralysis.^24^ Given the observed elevated frequency of T cells in the lymph node and the known neurotropism of ZIKV, we investigated T cells and their activation status in the brains of mice that were topically treated with VNAM. At day 7 pi, the time that neurological complications coincide with high virus loads in the brain, we observed a significantly increased CD69-expression on both CD8+ and CD4+ T cells compared to control mice (Fig. 7A-B). These data indicate more activated T cell populations in the brain, which could lead to extensive inflammation and tissue damage and in this way exacerbate neurological complications. These results thus indicate that T-cell mediated neuropathogenesis could be a key factor driving the accelerated disease progression and reduced survival observed after VNAM treatment.

As previous studies highlighted upregulation of IFN-stimulating genes after VNAM treatment^20^, we conducted RNA sequencing on footpad skin tissue to determine differences in gene expression between VNAM- and vehicle-treated mice. This analysis showed a significant downregulation of genes involved in muscle contraction functions (Fig. S9). Interestingly, a thin striated muscular layer, the panniculus carnosus, has been reported as a functional component of murine skin.^47^ Antibiotics have previously been linked to muscle atrophy, as seen with metronidazole-induced decrease in hind limb muscle weight and upregulation of genes involved in skeletal muscle neurogenic atrophy.^48^ Reports have also suggested that treatment with vancomycin and aminoglycosides such as neomycin induced myopathy and muscle weakness.^49,50^ Similarly, germ-free mice were reported with muscle atrophy and downregulation of genes associated with muscle growth, which were also downregulated in our study.^51^ Furthermore, ZIKV replication has already been detected in muscle cells and ZIKV-induced degeneration of muscle fibers has been observed.^16,52^ Hence, the combination of antibiotic treatment and ZIKV infection in our study could have exacerbated muscle atrophy. Further experiments are necessary to determine the extent to which muscle atrophy could have contributed as underlying mechanism in our model.

Finally, we aimed to elucidate whether the accelerated disease progression was directly caused by the local reduction of bacteria on the skin. To this end, we investigated if restoring the bacterial loads in the skin could rescue the antibiotic-induced phenotype. Bacterial CFUs in the skin were restored after the antibiotic treatment by inserting a 3 day period without treatment before ZIKV infection. This recolonization exceeded the bacterial CFUs in the skin of mice treated with control cream, potentially attributed to the less competitive environment promoting their growth more efficiently than in a stable microbial community (Fig. 8). Notably, the restoration of skin bacteria reversed the phenotype that was induced by the topical antibiotic treatment, delaying disease progression similar to what was observed in control mice. These data highlight a significant role of host skin bacteria in modulating ZIKV infection and disease outcome. In contrast, in the previously published study with oral antibiotic treatment in flavivirus-infected mice, susceptibility was not altered after oral microbiota transfer, suggesting a differential impact depending on the administration route of antibiotics and the microbial site.^19^

Achieving a comprehensive understanding of the interplay between skin bacteria and mosquito-borne viruses will require more detailed investigations utilizing approaches such as germ-free infection models or targeted manipulation of specific bacterial species (i.e. skin bacterial transplantations^53^). Such studies could pinpoint specific bacteria that modulate the susceptibility to ZIKV infection and clarify the specific underlying mechanisms. Integrating this knowledge with studies that identified bacteria species which attract or repel mosquitoes^11^, could yield valuable insights for the development of innovative preventive or therapeutic approaches based on microbiome manipulation. Additionally, more research will be essential regarding how these findings in immunodeficient AG129 mice can be effectively translated to humans. Considering that the skin serves as a common site of transmission for all mosquito-borne viruses, advancing our understanding of the interplay between these viruses and the host microbiome will be of utmost importance in counteracting not only ZIKV, but also other mosquito-borne viral diseases at early stages of the infection.

In conclusion, our study provided insights into the complex interplay between host bacteria, immune responses and viral pathogenesis and emphasized that potential influences of host bacteria should be considered in future mosquito-borne virus research. We identified antibiotic treatment of the skin as a risk factor for accelerated ZIKV disease progression and highlighted a substantial impact of host skin bacteria on ZIKV infection and dissemination.

## Acknowledgments.

This work was supported by a PhD fellowship granted to S.V. by the Research Foundation – Flanders (FWO) (11D5923N). We would like to thank the Rega Animalium staff for their support in animal experiments; Ji Ma, Geert Schoofs, Michael Schmid and Rafaela Vaz Sousa Pereira for their guidance to set up the flow cytometry experiments; Carolien De Keyzer, Elke Maas and Lindsey Bervoets for their help in animal experiments; Tina Van Buyten, Niels Cremers, Nanci Fereirra, Xin Zhang, Lara Kelchtermans, Elias Broeckhoven, Jana Meerkens, Stijn Verschoren, Katrien Trappeniers and Julia Zajac for their assistance in treatment of mice; Jasper Rymenants, Tina Van Buyten, Sadia Shaukat and Kristien Minner for their help in flow cytometry experiments; Joyce van Bree for technical support with DNA extractions; Elizabeth Grice and Julie Segre for their advice on bacterial DNA extractions from skin samples; Raul Yhossef Tito Tadeo for his support to set up 16S qPCR; Vanessa Brys and Annelien Verfaillie for the RNA sequencing of skin samples; Alvaro Cortes Calabuig for analyzing RNA sequencing data; Lanjiao Wang for sharing the 16S sequencing analysis pipeline; Inge Kortekaas for providing advice on skin flow cytometry; Katrien Lagrou for providing bacterial isolates; An Delang for preparing the antibiotic cream.

## Author contributions

Conceptualization, L.L., S.V. and L.D.; Methodology, L.L., S.V., S.J., R.A., Y.A.A., B.M.D., L.Y., and L.D.; Former Analysis, L.L., S.V., S.J., Y.A.A., B.M.D., C.C., L.Y.; Investigation, L.L., S.V., S.J., R.A., J.R., C.C., L.Y.; Resources, J.R., C.C., L.Y. L.D.; Writing – Original Draft, L.L., S.V., L.D.; Writing – Review & Editing, S.V., B.M.D., C.C., L.Y., L.D.; Visualization, L.L., S.V.; Supervision, L.D.; Project Administration, L.D.; Funding Acquisition, L.D.

## Supplemental information

Document S1. Figures S1-S9 and Table S1.

## Methods

### Contact for reagent and resource sharing

Further information and requests for resources and reagents should be directed to and will be fulfilled by the Lead Contact, Leen Delang.

### Cells and virus

Vero E6 cells (ATCC CRL-1586) were cultured in minimal essential medium (MEM, Gibco) supplemented with 10% fetal bovine serum (FBS, Gibco), 1% L-glutamine (Gibco), 1% sodium bicarbonate (Gibco) and 1% non-essential amino acids (NEAA; Gibco). Baby hamster kidney (BHK-21) cells (ATCC CCL-10) were cultured in Dulbecco’s Modified Eagle medium (DMEM, Gibco) supplemented with 10% FBS. Mammalian cell cultures were maintained at 37°C, 5% CO_2_ and 95%-99% relative humidity. *Aedes albopictus*-derived C6/36 cells (ATCC CRL-1660) were cultured in Leibovitz L-15 medium (Gibco) supplemented with 10% FBS, 1% NEAA, 1% 4-(2-hydroxyethyl)-1-piperazineethanesulfonic acid (HEPES; Gibco) and maintained at 28°C in the absence of CO_2_. For virus propagation and assays, similar cell culture media supplemented with 2% FBS (assay medium) were used.

The ZIKV PRVABC59 strain was acquired from the World Reference Center for Emerging Viruses and Arboviruses at the University of Texas Medical Branch (Genbank KU501215.1). Virus stocks were propagated in C6/36 cells and stored at -80°C. Viral titers were determined by endpoint titration on Vero E6 cells and by plaque assay on BHK-21 cells.

### Antibiotics

An oral antibiotic cocktail, consisting of vancomycin (0.35 g/L; Cayman Chemical), neomycin (1 g/L; Sigma-Aldrich), ampicillin (1 g/L; Sigma-Aldrich) and metronidazole (1g/L; Cayman Chemical) was prepared by dissolving compounds in water supplemented with 2.5% sucrose (Biofresh). The topical antibiotic cream containing vancomycin (0.5 g), neomycin (0.5 g), ampicillin (0.5 g), metronidazole (0.5 g) (VNAM), paraffin (15 g) and vaseline (33 g), and the control cream (containing paraffin (15 g) and vaseline (35 g)) were prepared by Apotheek Delang (Belgium). Bactericidal activity of the cream was visually confirmed by a disk-diffusion antibiotic sensitivity test. Liquid cultures of *Staphylococcus xylosus* were grown on BHI agar (Sigma) plates. Paper disks containing VNAM or control cream were placed on top of the agar surface and plates were incubated overnight at 37°C. The presence or absence of a zone of inhibition was visually checked the next day.

### Mice

Mouse experiments were performed with the approval and under the guidelines of the Ethical Committee of the University of Leuven (licenses P071/2019 and M020/2020). In-house-bred, female AG129 mice (deficient in both IFN-α/β and IFN-γ receptors) were maintained under specific pathogen-free (SPF) conditions at the Animal Facility of the Rega Institute for Medical Research, Leuven, Belgium. Mice were housed in individually ventilated isolator cages (IsoCage N Biocontainment System, Tecniplast) under standard conditions (18-23°C, 14 h:10 h light:dark cycle and 40-60% relative humidity) with cage enrichment and access to food and water ad libitum.

### Compound cytotoxicity and antiviral CPE reduction assay

To test the cytotoxicity and potential antiviral efficacy of the antibiotics, Vero E6 cells were seeded (10^4^ cells/well) in 96-well plates and incubated overnight (37°C, 5% CO_2_). The next day, 1:2 dilution series of the antibiotics were prepared in 2% assay medium on the cells. At day 7 post infection (pi), the 50% cytotoxic/cytostatic concentration (CC_50_), was determined by colorimetric read out using the MTS/PMS method (Promega). In parallel, antiviral activity of the antibiotics was determined by adding 1:2 serial dilutions of the antibiotics to the cells, followed by infection with ZIKV (MOI 0.1). At day 7 pi, intracellular viral RNA was isolated by incubating cells with cell-to-cDNA lysis buffer (Ambion) at room temperature for 15 min and at 75°C for another 15 min, after which viral genome copies were quantified by qRT-PCR as described below.

### Survival studies in ZIKV-infected AG129 mice

Two survival studies were conducted to assess the impact of prophylactic antibiotic treatment on ZIKV infection. In the first study, 12- to 14-week-old female AG129 mice were infected with 1000 plaque forming units (PFU) of ZIKV; in the second study 9- to 12-week-old female AG129 mice were infected with 100 PFU (n=8 per group). Mice were prophylactically treated by topical application of an antibiotic cream containing VNAM on the left footpad skin, twice daily for 7 days (as previously described).^54^ The positive control group received the VNAM cocktail orally via the drinking water supplemented with sucrose with ad libitum access for 7 days.^19^ For the negative control group, vehicle cream was applied on the footpad skin. All mice were transferred to clean cages on the first, fourth and final day of treatment. At the end of treatment, mice were intraperitoneally anesthetized using a mixture of atropine (0.4 µl/g; Sterop), ketamine (0.4 µl/g; EuroVet), xylazine (0.8 µl/g; V.M.D.) and water (0.4 µl/g). Subsequently, mice were subcutaneously infected with ZIKV PRVABC59 (100 or 1000 PFU) in the left footpad. Submandibular blood was collected at day 3, 5 and 7 pi and serum was obtained after centrifugation (10 000 rpm, 10 min, 4°C). Mice were observed daily for weight loss and the development of ZIKV-induced disease. In case of weight loss exceeding 20% of their initial weight or in case of presence of severe disease signs (a.o. paralysis of the hindlimbs, conjunctivitis, hunched back) mice were euthanized by intraperitoneal injection with Dolethal (200 mg/ml sodium pentobarbital, Vétoquinol). Brain tissue was collected in RLT lysis buffer (Qiagen) following transcardial perfusion with PBS (Gibco). For the bacteria rescue study, a similar survival study was performed with 9-to-12-week-old female AG129 mice (n=8 per group). Here, an additional group of mice was included that received treatment with VNAM cream but that were infected (ZIKV, 1000 PFU) 3 days after the final topical treatment.

### RNA isolation and ZIKV qRT-PCR

Viral RNA was isolated from serum samples with the NucleoSpin RNA virus kit (Machery-Nagel) according to the manufacturer’s protocol. Tissue samples were homogenized in Precellys tubes containing 2.8 mm zirconium oxide beads (Bertin Instruments) and lysis buffer using an automated homogenizer (Precellys24, Bertin Instruments) (program: 2 cycles of 20 s at 7600 rpm with a 20 s interval). Homogenates were centrifuged (15 000 rpm, 10 min, 4°C) and total RNA was extracted from the supernatant using the RNeasy Mini Kit (Qiagen) or the E.Z.N.A Total RNA kit I (Omega-Biotek), according to the manufacturer’s protocols.

To quantify viral RNA, a one-step, quantitative RT-PCR was performed in a total volume of 2511μL, containing 13.9411μL RNase free water (Promega), 6.2511μL master mix (Eurogentec), 0.37511μL of forward primer (511-CCGCTGCCCAACACAAG-311; final concentration: 900 nM), 0.37511μL of reverse primer (511-CCACTAACGTTCTTTTGCAGACAT-3’; final concentration: 90011nM), 111μL of probe (511-FAM-AGCCTACCT-ZEN-TGACAAGCAATCAGACACTCAA-IABkFQ-311; final concentration: 20011nM), 0.062511μL reverse transcriptase (Eurogentec), and 311μL RNA sample. The qRT-PCR was performed with the Applied Biosystems 7500 fast real-time PCR system using the following conditions: 30 min at 48°C, 10 min at 95°C, followed by 40 cycles of 15 s at 95°C and 111min at 60°C. For quantification, standard curves were generated each run using 10-fold dilutions of a synthesized gene block coding for the ZIKV E protein (IDT). Limits of detection (LODs) were determined as the lowest viral loads that could be detected by the qRT-PCR assay in 95% of experiments^55^, taking into account the buffer volumes and tissue weights.

### Endpoint virus titration

Mouse tissue samples were collected in Precellys tubes containing 2% assay medium and were homogenized in two cycles at 7600 rpm with a 20-s interval. After centrifugation (15 00011rpm, 1011min, 4°C), supernatant was collected and infectious virus was quantified by endpoint titrations. To this end, Vero E6 cells were seeded in 96-well plates at a density of 10^4^ cells/well and allowed to adhere overnight (37°C, 5% CO_2_). The next day, 3 parallel 10-fold serial dilutions of the tissue homogenates or serum samples were prepared on the cells. After 711days, the cells were examined microscopically for virus-induced cytopathogenic effect (CPE) and scored comparing to the uninfected controls. The TCID_50_/mL (tissue culture infectious dose 50%) was calculated using the method of Reed and Muench^56^ and is defined as the virus dose that would infect 50% of the cell cultures. Limits of quantification (LOQs) were determined as the lowest viral loads that could be quantified using this method, taking into account buffer volumes and tissue weights.

### DNA isolation of feces and skin samples

Before the start and at the end of the VNAM treatment, fresh feces samples were collected in 2 mL Precellys tubes containing PBS at a ratio of 100 µL per 10 mg of sample. Fecal pellets were homogenized in three cycles at 6800 rpm with a 10-s interval. After centrifugation (10 00011rpm, 511min), supernatant was collected and total DNA was isolated using the QIAamp DNA mini kit according to the manufacturer’s protocol.

Additionally, sterile foam swabs (Puritan) were used to swab the footpad skin extensively for at least 30 sec and collected in PowerBead Pro Tubes (Qiagen). DNA was isolated using the DNeasy PowerSoil Pro kit (Qiagen) according to the manufacturer’s protocol.

### 16S qRT-PCR

Bacterial loads in feces samples were quantified by 16S qRT-PCR. The reaction mixture (2011μL) contained 4.67 μL nuclease free water (Bio-Rad), 10 μL SYBR green master mix (Bio-Rad), 0.67 μL of forward primer (511-TCCTACGGGAGGCAGCAGT-311)58, 0.6711μL of reverse primer (511-GGACTACCAGGGTATCTAATCCTGTT-311; final concentration of each primer: 2 µM), and 4 μL of isolated DNA. The qRT-PCR was performed with the Applied Biosystems 7500 fast real-time PCR system using the following conditions: 10 min at 95°C, followed by 40 cycles of 15 s at 95°C and 111min at 60°C. For quantification, standard curves were generated each run using 10-fold dilutions of a synthesized gene block containing a fragment of the 16S rRNA gene (IDT).

### 16S rRNA gene sequencing

For bacteria community profiling in feces samples, DNA was isolated as described above and sent to Macrogen for 16S metagenomic sequencing using the Illumina sequencing-by-synthesis technology. FASTQ files were provided by Macrogen and analysis was performed using R 4.3.1. Read quality control, filtering and trimming was performed by implementing the DADA2 package (https://benjjneb.github.io/dada2/tutorial.html). Taxonomy was assigned using the SILVA version 132 as reference taxonomy database. Next, the phyloseq package was used to calculate alpha diversity measures by Shannon and Simpson diversity indices.

For sequencing of skin bacterial communities, DNA was isolated from skin swabs as described above. Quality of DNA isolations was verified on a 1.5% agarose gel (Life Technologies) and concentrations were determined using a QuantiFluor® dsDNA kit with a detection limit of 50 pg/ml and a sensitivity range of 0.01 to 200 ng/µl. For sequencing of the 16S rRNA gene, a 16S library was prepared and next generation sequencing was carried out as described previously.^57^ The V4 region of the 16S rRNA gene was amplified using the 515F (GTGYCAGCMGCCGCGGTAA) and the 806R (GGACTACNVGGGTWTCTAAT) primers, which were modified Illumina adapters and adapters for directional sequencing. Next, the Illumina MiSeq platform (Illumina) was used for sequencing, following manufacturer’s guidelines. Amplicon data was processed using mothur software following guidelines as outlined in before.^59^ After assembly of contigs and exclusion of ambiguous base calls, sequences were aligned to the silva_seed database and trimmed as described previously.^61^ Next, chimeras were removed by vsearch. Sequences were classified based on a naïve Bayesian classifier against the Ribosomal Database Project (RDP) 16S rRNA gene training set (v 16), using an 85% cut-off for the pseudobootstrap confidence score. Sequencies identified as Archaea, Eukaryota, Chloroplasts, unknown, or Mitochondria at the kingdom level were excluded. Remaining data were clustered into OTUs at a 3% dissimilarity level, resulting in an OTU table. Further analysis was performed in R, where the OTU table generated by the mothur pipeline was imported. OTUs that appeared only once across all samples (singletons) were excluded from the analysis, after which rarefaction curves were plotted to assess sequencing depth. Sequences were classified using the least common ancestor (LCA) method based on the various taxonomies available on the SINA server (v1.2.11).^62^ Data was rarefied to 6734 sequences per sample and alpha diversity for each sample, as well as inter-sample distances were calculated using QIIME.^64^ Alpha diversity was measured by Shannon index, and beta diversity was measured by Canberra, Jaccard and Bray-Curtis metrics followed by principal coordinate analysis (PCoA) using QIIME and visualized in Emperor.^67^

### Culturing of skin bacteria

To quantify culturable bacteria of the footpad skin of mice, complete footpad skin samples were collected using sterile forceps and transferred to a 2 mL Precellys tube containing PBS. Samples were then homogenized in two cycles at 7600 rpm with a 20-s interval and shortly centrifuged. Next, 50 µl of undiluted or 10-fold diluted supernatant was streaked onto blood agar plates (provided by UZ Leuven, Laboratory Medicine) whereafter plates were incubated at 37°C overnight. Following incubation, colony forming units (CFUs) were determined.

### Replication kinetics of ZIKV in AG129 mice

ZIKV *in vivo* replication kinetics were determined in 11- to 14-week-old female AG129 mice (n=5 per group) (first study) and in 9- to 13-week old female AG129 mice (n=8 per group) (second study) infected with ZIKV. Mice used in the first study were not treated prior to ZIKV infection, whereas both untreated mice as mice treated with control or VNAM cream on the skin inoculation site were used in the second study. Mice were topically treated as described earlier and transferred to clean cages every 3 days. After the final treatment, they were anaesthetized and subcutaneously infected in the footpad with ZIKV PRVABC59 (1000 PFU). At different time points post-infection (first study: 2 h, 8 h, 16 h, 24 h, 48 h and 72 h pi; second study: 16 h, 48 h, 7 d, 9 d pi), mice were sacrificed as described earlier. Blood was collected via cardiac puncture, after which transcardial perfusion was performed using PBS. Half of footpad skin, spleen, brain and spinal cord were collected in 2% assay medium to perform endpoint titration. The other half of all tissues and the complete draining inguinal lymph node were collected in TRK lysis buffer (Omega-Biotek) to perform RNA isolation and qRT-PCR as described above. Blood was centrifuged (10 000 rpm, 10 min, 4°C) to obtain serum. Serum and tissues were stored at −80°C until further processing.

### Flow cytometry of skin, lymph node and brain tissue

Female 9 to 11-week-old AG129 mice (n=10 per group) were treated with control or VNAM cream and infected with ZIKV (PRVABC59, 1000 PFU) in the left footpad, as described earlier. Before the infection and at both 16 h and 48 h pi, footpad skin and draining inguinal lymph nodes were collected on ice in Hank’s Balanced Salt Solution without calcium/magnesium (HBSS, Gibco) and Roswell Park Memorial Institute (RPMI, Gibco) medium, respectively. In another study using similar conditions, draining inguinal lymph nodes and brain tissue were collected at day 7 or day 9 pi in RPMI after transcardiac perfusion with ice cold saline (Gibco). All tissues were kept on ice until dissociation.

Footpad skin was cut into triangular pieces and digested with Dispase II (Roche; 5 mg/ml in HBSS) for 1 h at 37°C^58^. Next, epidermis and dermis were separated. The dermis was enzymatically digested in a 6-well plate (BD Falcon) containing a mixture of collagenase I (Gibco; end concentration of 1.6 mg/ml) and DNase I (Roche; end concentration of 0.1 mg/ml) in DMEM. The dermis was cut into small pieces and incubated at 37°C for 90 min. Every 30 min, the tissue suspension was mixed by pipetting. After incubation, dermis samples were filtered through a 70 µm cell strainer (VWR) and were centrifuged at 400 g for 10 min at 4°C. Samples were washed with FACS buffer (PBS supplemented with 2% FBS and 2 mM EDTA (Gibco)) and were centrifuged at 400 g for 5 min at 4°C. To obtain single cells from the epidermis, the samples were transferred to microcentrifuge tubes (Eppendorf) containing trypsin-EDTA (Gibco; 0.05% in HBSS). The epidermis was cut into small pieces and was incubated at 37°C for 5 min. Samples were shaken and were again incubated at 37°C for 5 min. Next, trypsin neutralization solution (5% FBS in HBSS) was added to stop the digestion. Samples were filtered through a 70 µm cell strainer and were centrifuged (400 g, 10 min, 4°C). Epidermal cells were resuspended, added to dermal cells and complete cell suspensions were transferred to a round bottom 96-well plate (Corning). After centrifugation (1500 rpm, 5 min, 4°C), cells were resuspended in PBS. Next, cells were stained for 15 min with 1:2000 Zombie aqua viability dye (BV510; Biolegend) and 1:200 FcR blocking reagent (Miltenyi Biotec) in PBS. Afterwards, cells were stained with the following antibodies (Biolegend) for 25 min: cluster of differentiation (CD) 45 (Alexa Fluor 700, 1:100), CD3 (APC Fire 750, 1:200), CD19 (PE-Cy5, 1:200), CD64 (PE, 1:200), CD11b (BUV395, 1:200), CD11c (BV421, 1:200), major histocompatibility complex class II (MHC-II) (BV786, 1:400) and Ly6C (PerCP-Cy5.5, 1:300). Next, cells were fixed and permeabilized with Fix/Perm buffer (BD Biosciences) for 30 min and were stained with the anti-CD207 antibody (APC, 1:200; Biolegend) for 30 min. Finally, cells were washed with permeabilization buffer (BD Biosciences), resuspended in FACS buffer and filtered through a 70 µm cell strainer. Flow cytometry data were recorded with LSRFortessa cell analyzer (BD Biosciences) with 405, 488, 561 and 643 nm laser excitation lines and analyzed using FlowJo 10.8.1 software (TreeStar).

After collection of inguinal lymph nodes, single-cell suspensions were prepared by mechanical dissociation, followed by filtration through 70 µm mesh. Samples were centrifuged (400 g, 5 min, 4°C), whereafter cells were resuspended in FACS buffer. Non-specific binding was blocked using 2.4G2 supernatant and dead cells were labeled by fixable viability dye eFluor 780 (APC-H7, 1:2000; ThermoFisher). Cellular phenotypes were assessed using a flow cytometry panel containing markers to identify cell types. Data were acquired on a BD FACSymphony, with a panel covering CD45 (APC, 1:800; BD), CD4 (BUV805, 1:200; BD), CD8α (BYG584, 1:200; BD), CD19 (BV786, 1:100; BioLegend), natural killer cell lectin-like receptor NK1.1 (Pe-Cy5.5, 1:600; BD), T cell receptor (TCR) β (FITC, 1:600; BD), Ly6C (BYG790, 1:200; ThermoFisher), Ly6G (BV570, 1:600; ThermoFisher), CD11c (BV421, 1:400; BD) and SiglecF (BUV615, 1:400; BioLegend). Flow cytometry data collection was performed using FACSDIVA version 8.0.2 (BD). Flow data was analyzed using FlowJo version 10.7.

Brain tissues were finely minced and mechanically dissociated by pipetting, followed by enzymatic digestion using collagenase (0.2 mg/ml) and DNase I (10 mg/ml) in a total volume of 5 mL completed with RPMI. Samples were incubated shaking at 37°C for 30 min and again mechanically dissociated by pipetting. Resulting suspensions were filtered through 70 µm cell strainers with HBSS. After centrifugation of cells (10 min, 300 g, 4°C; max acceleration, half brake), pellets were resuspended in DMEM containing 30% isotonic Percoll (SIP; Sigma-Aldrich) in 10X PBS (Gibco). Next, 70% SIP in HBSS was carefully pipetted underneath to create a density gradient and samples were centrifuged (20 min, 500 g, RT; low acceleration, no brake). Hereafter, the interphase layer was transferred and diluted in HBSS to centrifuge again 10 min, 300g, 4°C; max acceleration, max brake). The cell pellet was then resuspended in FACS buffer and stained in a round bottom 96-well plate. Cells were stained for 15 min with 1:2000 Zombie aqua viability dye (BV510) and 1:200 FcR blocking reagent in PBS. Afterwards, cells were stained with the following antibodies (Biolegend) for 25 min: CD3 (PerCP, 1:100), CD4 (BV421, 1:1000), CD8 (Spark UV, 1:1500), CD44 (BV605, 1:200), CD62L (K Blue 520, 1:400), CD69 (BV786, 1:100). Finally, cells were resuspended in FACS buffer. Flow cytometry data were recorded with LSRFortessa cell analyzer (BD Biosciences) with 405, 488, 561 and 643 nm laser excitation lines and analyzed using FlowJo 10.8.1 software.

### RNA sequencing of skin

The footpad skin of 13 to 15-week-old female AG129 mice (n=5 per group) were treated with VNAM or control cream as described earlier. Mice were transferred to clean cages every 3 days. After treatment, mice were sacrificed and the left footpad skin was collected in RNA later® storage reagent. RNA was isolated with the E.Z.N.A Total RNA kit I as described above. RNA samples were sent to the Genomics Core (Center for Human Genetics) of KU Leuven/UZ Leuven. The quality of RNA was determined using the Bioanalyzer (Agilent) and concentration and purity was measured with Nanodrop (Thermo Fisher Scientific). Libraries were constructed using the TruSeq stranded mRNA preparation kit (Illumina). The obtained libraries were quantified using the Qubit RNA High Sensitivity assay kit (Thermo Fisher Scientific) and the average library size was determined with the Fragment Analyzer (Agilent). Library quantification was performed by qPCR analysis using the Kapa library quantification kit (Roche), whereafter equimolar pools were prepared. After denaturation, library pools were sequenced using single-read sequencing on the HISeq 4000 (Illumina).

Quality control of raw reads was performed with FastQC v0.11.7 (https://www.bioinformatics.babraham.ac.uk/projects/fastqc). Adapters were filtered with ea-utils fastq-mcf v1.05 (https://github.com/ExpressionAnalysis/ea-utils). Splice-aware alignment was performed with HISAT2^60^ against the mouse reference genome mm10 using the default parameters. Reads mapping to multiple loci in the reference genome were discarded. Resulting BAM alignment files were handled with Samtools v1.5.^63^ Quantification of reads per gene was performed with HT-seq Count v0.10.0, Python v2.7.14.^65^ Count-based differential expression analysis was done with R-based (The R Foundation for Statistical Computing, Vienna, Austria) Bioconductor package DESeq2.^66^ Reported p-values were adjusted for multiple testing with the Benjamini-Hochberg procedure, which controls false discovery rate (FDR). Gene ontology analysis was performed with the Gene ontology enrichment analysis and visualization tool (http://cbl-gorilla.cs.technion.ac.il/) and pathway analysis was performed using the Ingenuity Pathway Analysis tool (Qiagen).

### Statistical analysis

All data were analyzed using GraphPad Prism 8.3.1. Significant differences in survival rate were analyzed using the log-rank (Mantel-Cox) test. Significant differences in viral loads in tissues and bacterial loads in feces and flow cytometry experiments at different time points were assessed using the two-way repeated measures ANOVA with Sidak’s correction. All other results were analyzed using the non-parametric Kruskal-Wallis with Dunn’s multiple comparisons test (comparing 3 independent groups) or Mann-Whitney U test (comparing 2 independent groups). Statistical significance threshold was assessed at p values < 0.05. Statistical details are described in the figure legends.

## References

1. Pierson, T.C., and Diamond, M.S. The continued threat of emerging flaviviruses. 10.1038/s41564-020-0714-0.

2. Pardy, R.D., and Richer, M.J. (2019). Zika Virus Pathogenesis: From Early Case Reports to Epidemics. Viruses 11. 10.3390/V11100886.

3. Hills, S.L., Fischer, M., and Petersen, L.R. (2017). Epidemiology of Zika Virus Infection. J Infect Dis 216, S868–S874. 10.1093/INFDIS/JIX434.

4. Regional Zika Epidemiological Update (Americas) August 25, 2017 - PAHO/WHO | Pan American Health Organization https://www.paho.org/en/regional-zika-epidemiological-update-americas-august-25-2017.

5. Liu, Z.Y., Shi, W.F., and Qin, C.F. (2019). The evolution of Zika virus from Asia to the Americas. Nat Rev Microbiol 17, 131–139. 10.1038/S41579-018-0134-9.

6. Krauer, F., Riesen, M., Reveiz, L., Oladapo, O.T., Martínez-Vega, R., Porgo, T. V., Haefliger, A., Broutet, N.J., and Low, N. (2017). Zika Virus Infection as a Cause of Congenital Brain Abnormalities and Guillain–Barré Syndrome: Systematic Review. PLoS Med 14, e1002203. 10.1371/JOURNAL.PMED.1002203.

7. Barbi, L., Coelho, A.V.C., Alencar, L.C.A. de, and Crovella, S. (2018). Prevalence of Guillain-Barré syndrome among Zika virus infected cases: a systematic review and meta-analysis. Braz J Infect Dis 22, 137–141. 10.1016/J.BJID.2018.02.005.

8. Pingen, M., Schmid, M.A., Harris, E., and McKimmie, C.S. (2017). Mosquito Biting Modulates Skin Response to Virus Infection. Trends Parasitol 33, 645–657. 10.1016/j.pt.2017.04.003.

9. Grice, E.A., and Segre, J.A. (2011). The skin microbiome. Nat Rev Microbiol 9, 244–253. 10.1038/nrmicro2537.

10. Byrd, A.L., Belkaid, Y., and Segre, J.A. (2018). The human skin microbiome. Nature Reviews Microbiology 2018 16:3 16, 143–155. 10.1038/nrmicro.2017.157.

11. Verhulst, N.O., Qiu, Y.T., Beijleveld, H., Maliepaard, C., Knights, D., Schulz, S., Berg-Lyons, D., Lauber, C.L., Verduijn, W., Haasnoot, G.W., et al. (2011). Composition of Human Skin Microbiota Affects Attractiveness to Malaria Mosquitoes. PLoS One 6, e28991. 10.1371/JOURNAL.PONE.0028991.

12. Zhang, H., Zhu, Y., Liu, Z., Peng, Y., Peng, W., Tong, L., Wang, J., Liu, Q., Wang, P., and Cheng, G. (2022). A volatile from the skin microbiota of flavivirus-infected hosts promotes mosquito attractiveness. Cell 185, 2510–2522.e16. 10.1016/J.CELL.2022.05.016.

13. Lana, L., Sofie, J., Rana, A., Sam, V., Suzanne, K., Pieter, B., Lieve, V.M., and Leen, D. (2022). Perturbation of alphavirus and flavivirus infectivity by components of the bacterial cell wall. J Virol 0, jvi.00060–22. 10.1128/jvi.00060-22.

14. Morrison, T.E. (2018). Animal Models for Chikungunya Virus and Zika Virus. Chikungunya and Zika Viruses: Global Emerging Health Threats, 317–346. 10.1016/B978-0-12-811865-8.00010-6.

15. Santos, F.M., Dias, R.S., de Oliveira, M.D., Costa, I.C.T.A., Fernandes, L. de S., Pessoa, C.R., da Matta, S.L.P., Costa, V.V., Souza, D.G., da Silva, C.C., et al. (2019). Animal model of arthritis and myositis induced by the Mayaro virus. PLoS Negl Trop Dis 13, e0007375. 10.1371/JOURNAL.PNTD.0007375.

16. Aliota, M.T., Caine, E.A., Walker, E.C., Larkin, K.E., Camacho, E., and Osorio, J.E. (2016). Characterization of Lethal Zika Virus Infection in AG129 Mice. PLoS Negl Trop Dis 10, e0004682. 10.1371/JOURNAL.PNTD.0004682.

17. Gardner, J., Anraku, I., Le, T.T., Larcher, T., Major, L., Roques, P., Schroder, W.A., Higgs, S., and Suhrbier, A. (2010). Chikungunya Virus Arthritis in Adult Wild-Type Mice. J Virol 84, 8021– 8032. 10.1128/JVI.02603-09/SUPPL_FILE/SUPPLEMENTAL_FIGURES.PDF.

18. SanMiguel, A.J., Meisel, J.S., Horwinski, J., Zheng, Q., and Grice, E.A. (2017). Topical Antimicrobial Treatments Can Elicit Shifts to Resident Skin Bacterial Communities and Reduce Colonization by Staphylococcus aureus Competitors. Antimicrob Agents Chemother 61, e00774–17. 10.1128/AAC.00774-17.

19. Thackray, L.B., Handley, S.A., Gorman, M.J., Poddar, S., Bagadia, P., Briseño, C.G., Theisen, D.J., Tan, Q., Hykes Jr., B.L., Lin, H., et al. (2018). Oral Antibiotic Treatment of Mice Exacerbates the Disease Severity of Multiple Flavivirus Infections. Cell Rep 22, 3440–3453.e6. 10.1016/j.celrep.2018.03.001.

20. Gopinath, S., Kim, M. V, Rakib, T., Wong, P.W., van Zandt, M., Barry, N.A., Kaisho, T., Goodman, A.L., and Iwasaki, A. (2018). Topical application of aminoglycoside antibiotics enhances host resistance to viral infections in a microbiota-independent manner. Nat Microbiol 3, 611–621. 10.1038/s41564-018-0138-2.

21. Li, C., Zu, S., Deng, Y.Q., Li, D., Parvatiyar, K., Quanquin, N., Shang, J., Sun, N., Su, J., Liu, Z., et al. (2019). Azithromycin Protects against Zika Virus Infection by Upregulating Virus-Induced Type I and III Interferon Responses. Antimicrob Agents Chemother 63. 10.1128/AAC.00394-19.

22. Yuan, J., Yu, J., Huang, Y., He, Z., Luo, J., Wu, Y., Zheng, Y., Wu, J., Zhu, X., Wang, H., et al. (2020). Antibiotic fidaxomicin is an RdRp inhibitor as a potential new therapeutic agent against Zika virus. BMC Med 18, 1–16. 10.1186/S12916-020-01663-1/FIGURES/6.

23. Winkler, E.S., Shrihari, S., Hykes Jr., B.L., Handley, S.A., Andhey, P.S., Huang, Y.-J.S., Swain, A., Droit, L., Chebrolu, K.K., Mack, M., et al. (2020). The Intestinal Microbiome Restricts Alphavirus Infection and Dissemination through a Bile Acid-Type I IFN Signaling Axis. Cell 182, 901–918.e18. 10.1016/j.cell.2020.06.029.

24. Jurado, K.A., Yockey, L.J., Wong, P.W., Lee, S., Huttner, A.J., and Iwasaki, A. (2017). Antiviral CD8 T cells induce Zika-virus-associated paralysis in mice. Nature Microbiology 2018 3:2 3, 141–147. 10.1038/s41564-017-0060-z.

25. Nazerai, L., Schøller, A.S., Bassi, M.R., Buus, S., Stryhn, A., Christensen, J.P., and Thomsen, A.R. (2020). Effector CD8 T Cell-Dependent Zika Virus Control in the CNS: A Matter of Time and Numbers. Front Immunol 11. 10.3389/FIMMU.2020.01977.

26. Manangeeswaran, M., Ireland, D.D.C., and Verthelyi, D. (2016). Zika (PRVABC59) Infection Is Associated with T cell Infiltration and Neurodegeneration in CNS of Immunocompetent Neonatal C57Bl/6 Mice. PLoS Pathog 12, e1006004. 10.1371/JOURNAL.PPAT.1006004.

27. Meisel, J.S., Sfyroera, G., Bartow-McKenney, C., Gimblet, C., Bugayev, J., Horwinski, J., Kim, B., Brestoff, J.R., Tyldsley, A.S., Zheng, Q., et al. (2018). Commensal microbiota modulate gene expression in the skin. Microbiome 6, 1–15. 10.1186/S40168-018-0404-9/FIGURES/5.

28. Belheouane, M., Vallier, M., Čepić, A., Chung, C.J., Ibrahim, S., and Baines, J.F. (2020). Assessing similarities and disparities in the skin microbiota between wild and laboratory populations of house mice. ISME J 14, 2367–2380. 10.1038/s41396-020-0690-7.

29. Tavakkol, Z., Samuelson, D., Delancey Pulcini, E., Underwood, R.A., Usui, M.L., William Costerton, J., James, G.A., Olerud, J.E., and Fleckman, P. (2010). Resident Bacterial Flora in the Skin of C57BL/6 Mice Housed under SPF Conditions.

30. Rubinstein, E., and Keynan, Y. (2014). Vancomycin revisited - 60 years later. Front Public Health 2. 10.3389/FPUBH.2014.00217.

31. Vardanyan, R.S., and Hruby, V.J. (2006). Antibiotics. Synthesis of Essential Drugs, 425–498. 10.1016/B978-044452166-8/50032-7.

32. Castle, S.S. (2007). Ampicillin. xPharm: The Comprehensive Pharmacology Reference, 1–6. 10.1016/B978-008055232-3.61227-9.

33. Kaushik, D., Mohan, M., Borade, D.M., and Swami, O.C. (2014). Ampicillin: rise fall and resurgence. J Clin Diagn Res 8. 10.7860/JCDR/2014/8777.4356.

34. Freeman, C.D., Klutman, N.E., and Lamp, K.C. (2012). Metronidazole. Drugs 1997 54:5 54, 679–708. 10.2165/00003495-199754050-00003.

35. Lubin, J.B., Green, J., Maddux, S., Denu, L., Duranova, T., Lanza, M., Wynosky-Dolfi, M., Flores, J.N., Grimes, L.P., Brodsky, I.E., et al. (2023). Arresting microbiome development limits immune system maturation and resistance to infection in mice. Cell Host Microbe 31, 554–570.e7. 10.1016/j.chom.2023.03.006.

36. Yang, J.H., Bhargava, P., McCloskey, D., Mao, N., Palsson, B.O., and Collins, J.J. (2017). Antibiotic-Induced Changes to the Host Metabolic Environment Inhibit Drug Efficacy and Alter Immune Function. Cell Host Microbe 22, 757–765.e3. 10.1016/j.chom.2017.10.020.

37. Soria, M.E., Cortón, M., Martínez-González, B., Lobo-Vega, R., Vázquez-Sirvent, L., López-Rodríguez, R., Almoguera, B., Mahillo, I., Mínguez, P., Herrero, A., et al. (2021). High SARS-CoV-2 viral load is associated with a worse clinical outcome of COVID-19 disease. Access Microbiol 3, 000259. 10.1099/ACMI.0.000259.

38. Reynoso, G. V., Gordon, D.N., Kalia, A., Aguilar, C.C., Malo, C.S., Aleshnick, M., Dowd, K.A., Cherry, C.R., Shannon, J.P., Vrba, S.M., et al. (2023). Zika virus spreads through infection of lymph node-resident macrophages. Cell Rep 42. 10.1016/J.CELREP.2023.112126.

39. McDonald, E.M., Anderson, J., Wilusz, J., Ebel, G.D., and Brault, A.C. (2020). Zika Virus Replication in Myeloid Cells during Acute Infection Is Vital to Viral Dissemination and Pathogenesis in a Mouse Model. J Virol 94. 10.1128/JVI.00838-20.

40. Fink, K., Ng, C., Nkenfou, C., Vasudevan, S.G., Van Rooijen, N., and Schul, W. (2009). Depletion of macrophages in mice results in higher dengue virus titers and highlights the role of macrophages for virus control. Eur J Immunol 39, 2809–2821. 10.1002/EJI.200939389.

41. Wang, Y.-T., Hattakam, S., Young, M.P., Regla-Nava, J.A., and Shresta, S. (2019). Monocytes and macrophages limit systemic infection and modulate the CD4 T cell response during Zika virus infection in mice. The Journal of Immunology 202, 140.5–140.5. 10.4049/JIMMUNOL.202.SUPP.140.5.

42. Zukor, K., Wang, H., Siddharthan, V., Julander, J.G., and Morrey, J.D. (2018). Zika virus-induced acute myelitis and motor deficits in adult interferon αβ/γ receptor knockout mice. J Neurovirol 24, 273–290. 10.1007/S13365-017-0595-Z/FIGURES/14.

43. Santara, S. Sen, Angela, A.C., Mulik, S., Ovies, C., Boulenouar, S., Strominger, J.L., and Lieberman, J. (2021). Decidual NK cells kill Zika virus-infected trophoblasts. Proc Natl Acad Sci U S A 118, e2115410118. 10.1073/PNAS.2115410118/SUPPL_FILE/PNAS.2115410118.SAPP.PDF.

44. Maucourant, C., Nonato Queiroz, G.A., Corneau, A., Leandro Gois, L., Meghraoui-Kheddar, A., Tarantino, N., Bandeira, A.C., Samri, A., Blanc, C., Yssel, H., et al. (2021). NK Cell Responses in Zika Virus Infection Are Biased towards Cytokine-Mediated Effector Functions. J Immunol 207, 1333–1343. 10.4049/JIMMUNOL.2001180.

45. Shabrish, S., Karnik, N., Gupta, V., Bhate, P., and Madkaikar, M. (2020). Impaired NK cell activation during acute dengue virus infection: A contributing factor to disease severity. Heliyon 6. 10.1016/J.HELIYON.2020.E04320.

46. Huang, H., Li, S., Zhang, Y., Han, X., Jia, B., Liu, H., Liu, D., Tan, S., Wang, Q., Bi, Y., et al. (2017). CD8+ T Cell Immune Response in Immunocompetent Mice during Zika Virus Infection. J Virol 91. 10.1128/JVI.00900-17.

47. Naldaiz-Gastesi, N., Bahri, O.A., López de Munain, A., McCullagh, K.J.A., and Izeta, A. (2018). The panniculus carnosus muscle: an evolutionary enigma at the intersection of distinct research fields. J Anat 233, 275–288. 10.1111/JOA.12840.

48. Manickam, R., Oh, H.Y.P., Tan, C.K., Paramalingam, E., and Wahli, W. (2018). Metronidazole Causes Skeletal Muscle Atrophy and Modulates Muscle Chronometabolism. International Journal of Molecular Sciences 2018, Vol. 19, Page 2418 19, 2418. 10.3390/IJMS19082418.

49. Al Alawi, K., Al Shinawy, M., Taqee, A., and Wali, Y. (2011). An Overlooked Complication of vancomycin induced acute flaccid paralysis in a child with acute leukemia: A case report. Oman Med J 26, 283–284. 10.5001/OMJ.2011.69.

50. Harnett, M.T., Chen, W., and Smith, S.M. (2009). Calcium-sensing receptor: A high-affinity presynaptic target for aminoglycoside-induced weakness. Neuropharmacology 57, 502–505. 10.1016/J.NEUROPHARM.2009.07.031.

51. Lahiri, S., Kim, H., Garcia-Perez, I., Reza, M.M., Martin, K.A., Kundu, P., Cox, L.M., Selkrig, J., Posma, J.M., Zhang, H., et al. (2019). The gut microbiota influences skeletal muscle mass and function in mice. Sci Transl Med 11, 5662. 10.1126/SCITRANSLMED.AAN5662/SUPPL_FILE/AAN5662_SM.PDF.

52. Rossi, S.L., Tesh, R.B., Azar, S.R., Muruato, A.E., Hanley, K.A., Auguste, A.J., Langsjoen, R.M., Paessler, S., Vasilakis, N., and Weaver, S.C. (2016). Characterization of a Novel Murine Model to Study Zika Virus. Am J Trop Med Hyg 94, 1362–1369. 10.4269/AJTMH.16-0111.

53. Callewaert, C., Knödlseder, N., Karoglan, A., Güell, M., and Paetzold, B. (2021). Skin microbiome transplantation and manipulation: Current state of the art. Comput Struct Biotechnol J 19, 624–631. 10.1016/J.CSBJ.2021.01.001.

54. SanMiguel, A.J., Meisel, J.S., Horwinski, J., Zheng, Q., and Grice, E.A. (2017). Topical antimicrobial treatments can elicit shifts to resident skin bacterial communities and reduce colonization by staphylococcus aureus competitors. Antimicrob Agents Chemother 61. 10.1128/AAC.00774-17/SUPPL_FILE/ZAC009176465S1.PDF.

55. Bustin, S.A., Benes, V., Garson, J.A., Hellemans, J., Huggett, J., Kubista, M., Mueller, R., Nolan, T., Pfaffl, M.W., Shipley, G.L., et al. (2009). The MIQE Guidelines: Minimum Information for Publication of Quantitative Real-Time PCR Experiments. Clin Chem 55, 611–622. 10.1373/CLINCHEM.2008.112797.

56. Reed, L.J., and Muench, H. (1938). A simple method of estimating fifty per cent endpoints. Am J Epidemiol 27, 493–497. 10.1093/OXFORDJOURNALS.AJE.A118408/2/27-3-493.PDF.GIF.

57. Vandeputte, D., Kathagen, G., D’Hoe, K., Vieira-Silva, S., Valles-Colomer, M., Sabino, J., Wang, J., Tito, R.Y., De Commer, L., Darzi, Y., et al. (2017). Quantitative microbiome profiling links gut community variation to microbial load. Nature 2017 551:7681 551, 507–511. 10.1038/nature24460.

58. Lou, F., Sun, Y., and Wang, H. (2020). Protocol for Flow Cytometric Detection of Immune Cell Infiltration in the Epidermis and Dermis of a Psoriasis Mouse Model. STAR Protoc 1, 100115. 10.1016/J.XPRO.2020.100115.

59. Kozich, J.J., Westcott, S.L., Baxter, N.T., Highlander, S.K., and Schloss, P.D. (2013). Development of a dual-index sequencing strategy and curation pipeline for analyzing amplicon sequence data on the MiSeq Illumina sequencing platform. Appl Environ Microbiol 79, 5112–5120. 10.1128/AEM.01043-13.

60. Kim, D., Paggi, J.M., Park, C., Bennett, C., and Salzberg, S.L. (2019). Graph-based genome alignment and genotyping with HISAT2 and HISAT-genotype. Nature Biotechnology s2019 37:8 37, 907–915. 10.1038/s41587-019-0201-4.

61. Quast, C., Pruesse, E., Yilmaz, P., Gerken, J., Schweer, T., Yarza, P., Peplies, J., and Glöckner, F.O. (2013). The SILVA ribosomal RNA gene database project: improved data processing and web-based tools. Nucleic Acids Res 41. 10.1093/NAR/GKS1219.

62. Pruesse, E., Peplies, J., and Glöckner, F.O. (2012). SINA: accurate high-throughput multiple sequence alignment of ribosomal RNA genes. Bioinformatics 28, 1823–1829. 10.1093/BIOINFORMATICS/BTS252.

63. Li, H., Handsaker, B., Wysoker, A., Fennell, T., Ruan, J., Homer, N., Marth, G., Abecasis, G., and Durbin, R. (2009). The Sequence Alignment/Map format and SAMtools. Bioinformatics 25, 2078–2079. 10.1093/BIOINFORMATICS/BTP352.

64. Kuczynski, J., Stombaugh, J., Walters, W.A., González, A., Caporaso, J.G., and Knight, R. (2012). Using QIIME to analyze 16S rRNA gene sequences from microbial communities. Curr Protoc Microbiol Chapter 1. 10.1002/9780471729259.MC01E05S27.

65. Anders, S., Pyl, P.T., and Huber, W. (2015). HTSeq—a Python framework to work with high-throughput sequencing data. Bioinformatics 31, 166–169. 10.1093/BIOINFORMATICS/BTU638.

66. Love, M.I., Huber, W., and Anders, S. (2014). Moderated estimation of fold change and dispersion for RNA-seq data with DESeq2. Genome Biol 15, 1–21. 10.1186/S13059-014-0550-8/FIGURES/9.

67. Vázquez-Baeza, Y., Pirrung, M., Gonzalez, A., and Knight, R. (2013). EMPeror: a tool for visualizing high-throughput microbial community data. Gigascience 2. 10.1186/2047-217X-2-16.

