## Supplemental information for "Depletion of skin bacteria by topical antibiotic treatment accelerates onset of Zika virus disease in mice"

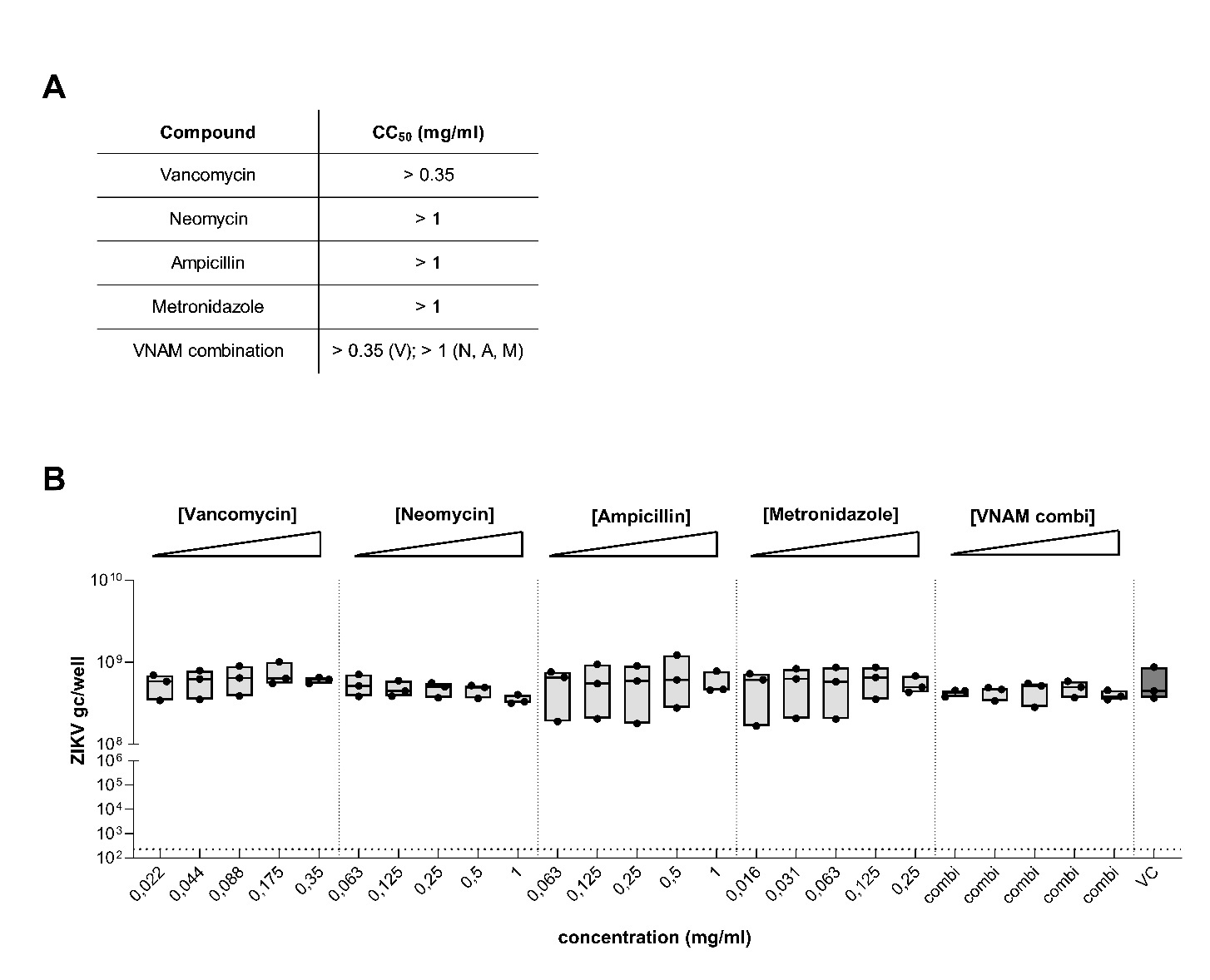
**Supplemental figures.**

**Fig. S1. VNAM antibiotics (single and combination) had no toxic or anti-ZIKV activity *in vitro*.**

(A) CC_50_ values of vancomycin, neomycin, ampicillin, metronidazole and the VNAM combination in VeroE6 cells, determined by MTS read out. (B) *In vitro* antiviral effect of vancomycin, neomycin, ampicillin, metronidazole and the VNAM combination determined in Vero E6 cells infected with ZIKV (MOI 0.01). ZIKV intracellular RNA (genome copies/well) was quantified at day 7 pi by qRT-PCR. Data from 3 independent experiments are represented as floating bars, showing the individual data points with solid lines indicating the median values. Dotted line indicates the LOD. CC50: cytotoxic concentration 50%; LOD: limit of detection; VC: virus control; gc: genome copies.

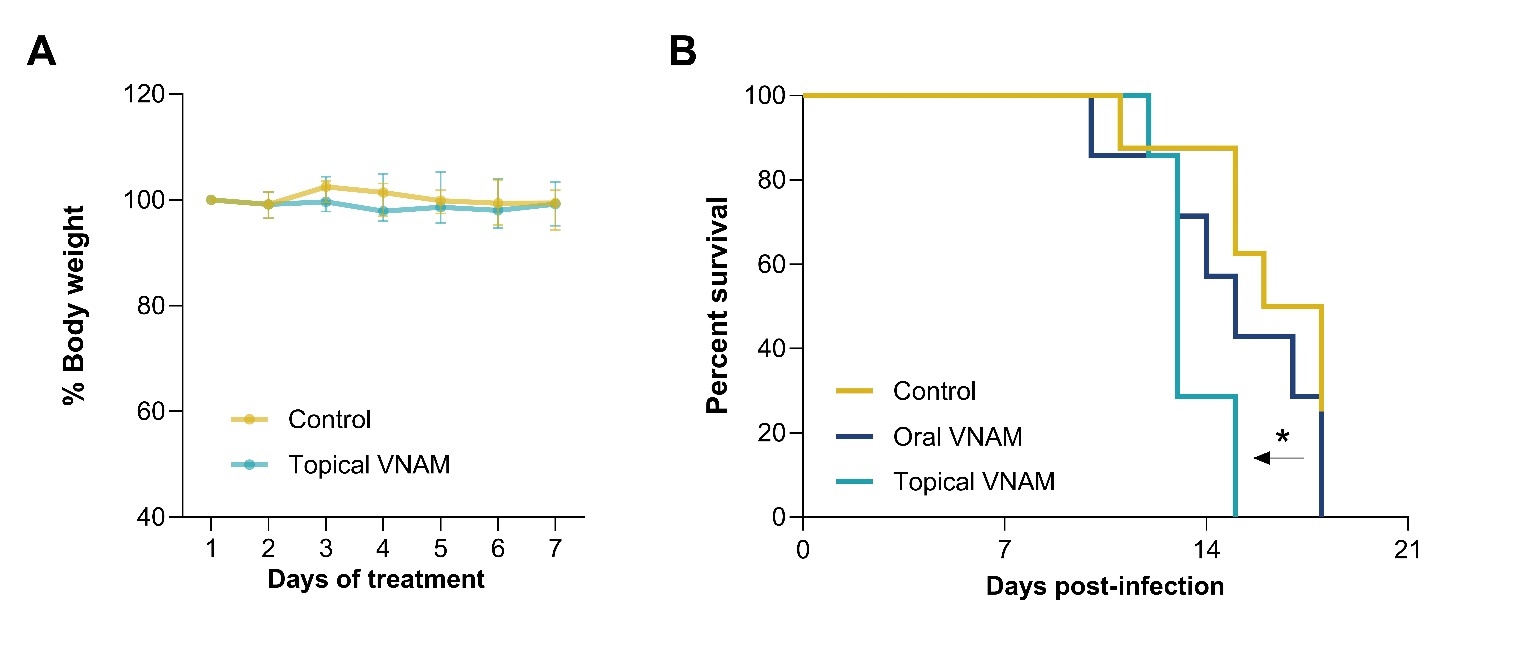

**Fig. S2. Antibiotic treatment accelerated disease progression in mice infected with a lower ZIKV inoculum.**

Mice were treated with an oral VNAM cocktail, a VNAM cream or a vehicle cream at the skin inoculation site (n=8 per group). Subsequently, mice were subcutaneously infected in the footpad with the ZIKV PRVABC59 strain (100 PFU). (A) Percentage of body weight of mice treated with VNAM and vehicle cream over the 7-day treatment period, compared to their initial body weight. (B) Kaplan-Meier survival curves after ZIKV infection. Median values of survival were calculated and a log-rank (Mantel-Cox) test was performed to assess statistically significant differences between survival curves (*, p<0.05). VNAM: vancomycin, neomycin, ampicillin, metronidazole.

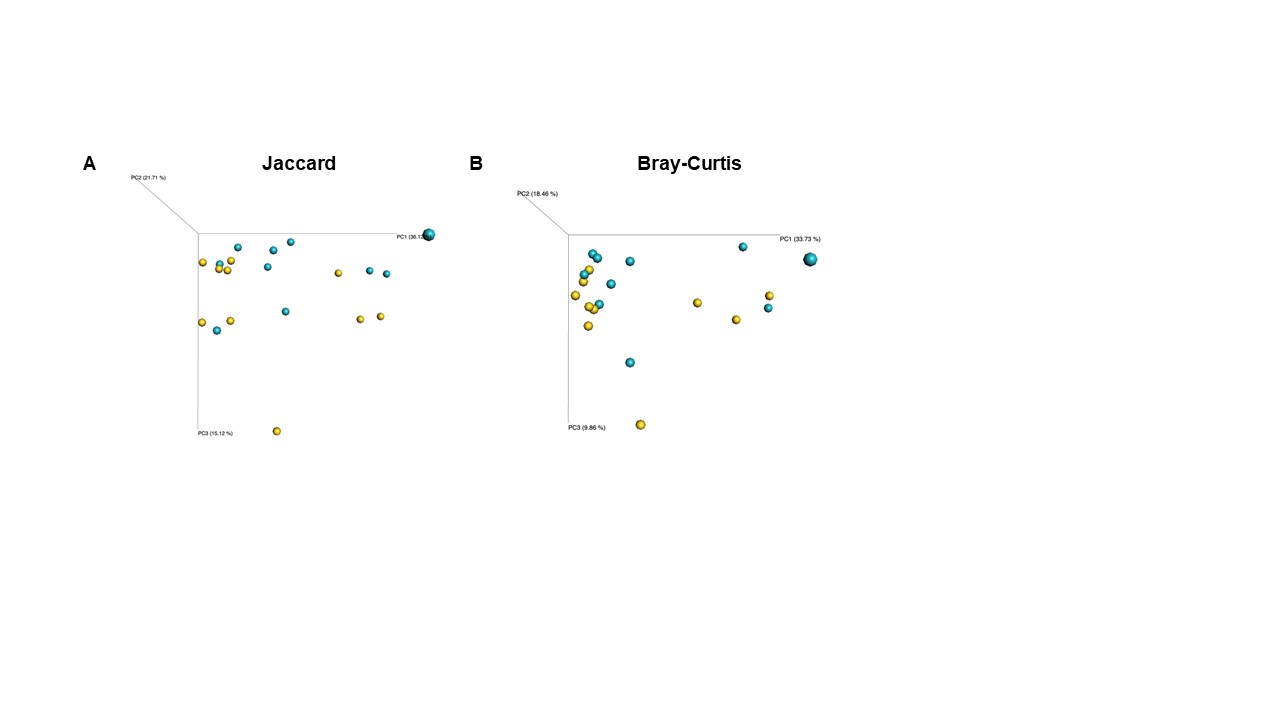

**Fig. S3. Topical antibiotic treatment induced a shift in bacterial community composition in the skin.**

Mice were orally or topically treated with VNAM or vehicle (n=10 per group) and skin swabs were collected after 7 days of treatment. Beta-diversity based PCoA plots using (A) Jaccard or (B) Bray-Curtis metrics of skin bacterial communities after 16S metagenomic sequencing of skin swabs. PC: principal component.

**Fig. S4. Replication kinetics of ZIKV at the skin inoculation site and in secondary organs.**

Mice were infected with ZIKV (1000 PFU) (n=5 per group). Tissues were collected and viral RNA was quantified by qRT-PCR at 2, 8, 16, 24, 48 and 72 h pi. Viral RNA in the (A) skin, (B) draining lymph node, (C) serum and (D) spleen. Each data point represents an individual mouse, with solid lines representing the median values. Statistical significance was assessed with the Mann-Whitney U test (*, p<0.05; **,
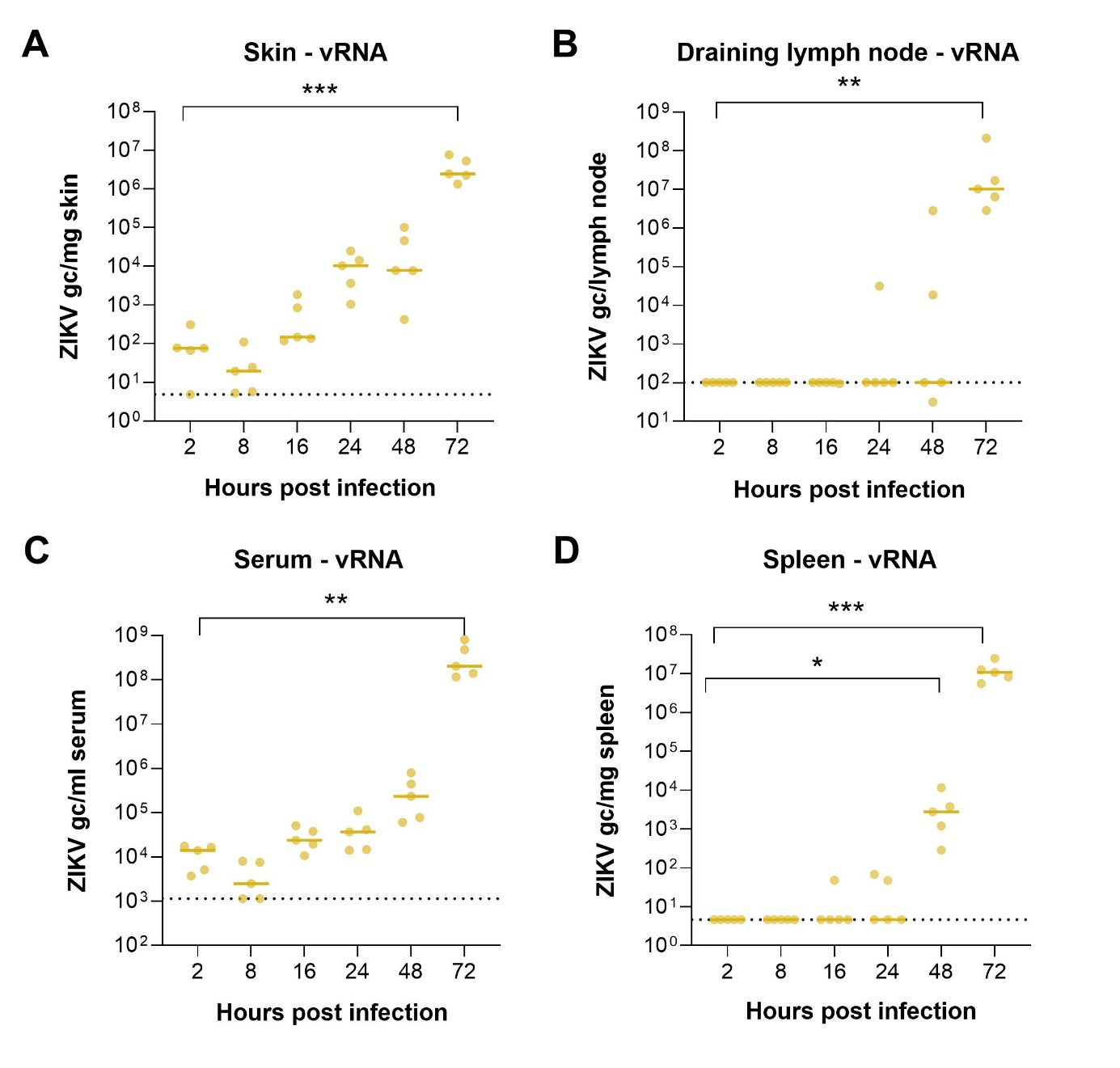
p<0.01; ***, p<0.001). The dotted lines represent the LOD. vRNA: viral RNA; gc: genome copies; LOD: limit of detection.

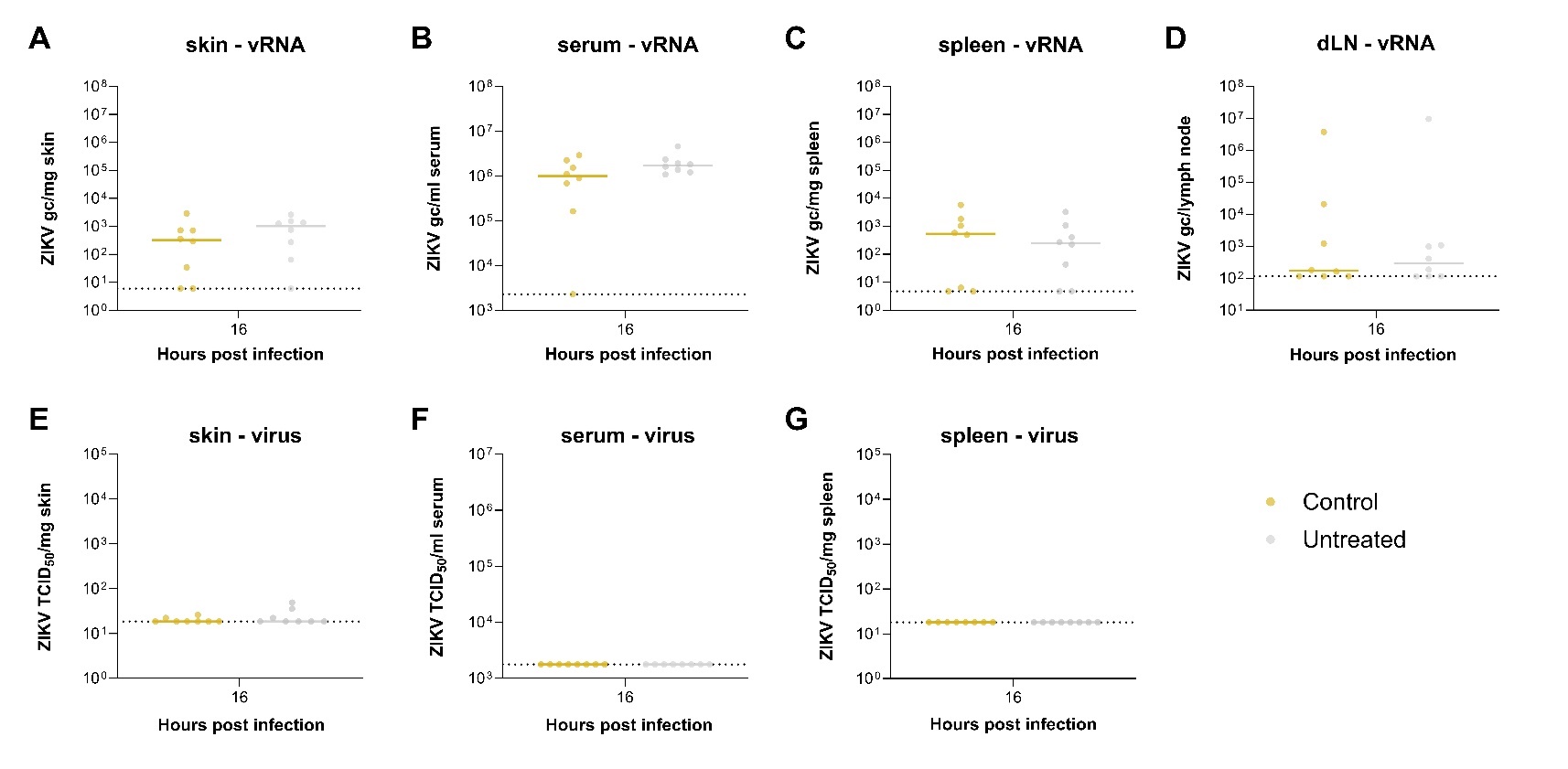

**Fig. S5. Control treatment did not influence virus replication.**

Mice were topically treated with vehicle or no treatment (n=8 per group) prior to infection with ZIKV (1000 PFU). Tissues were analyzed at 16 h pi. ZIKV RNA levels in (A) skin of the inoculation site, (B) serum, (C) spleen, (D) draining lymph node, as determined by qRT-PCR. TCID_50_ values in (E) skin of the inoculation site, (F) serum and (G) spleen, quantified by viral end-point titrations on Vero cells. Individual data points are shown, with solid lines representing the median value. Statistically significant differences between control and untreated mice were assessed with a two-way ANOVA with Sidak’s correction for multiple comparisons (ns, p>0.05). (A-D) Dotted lines represent the LOD. (E-G) Dotted lines represent the LOQ. vRNA: viral RNA; gc: genome copies; pi: post infection; TCID50: tissue culture infectious dose 50; LOD: limit of detection; LOQ: limit of quantification.

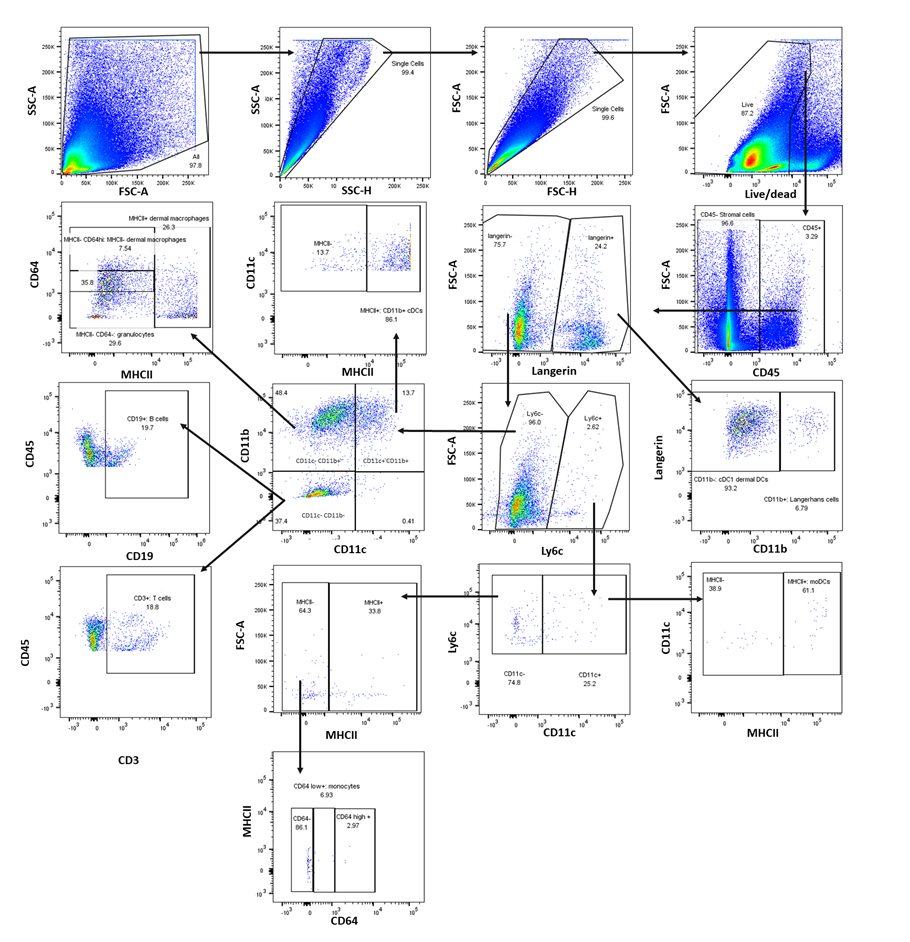

**Fig. S6. Gating strategy for the flow cytometric analysis of immune cells in the skin.**

Pseudocolor plots show the gating of skin cells based on surface markers (CD45, Ly6C, CD11b, CD11c, MHC-II, CD64, CD3, CD19) and an intracellular marker (CD207). Percentages indicate frequencies of cells of the parent population.

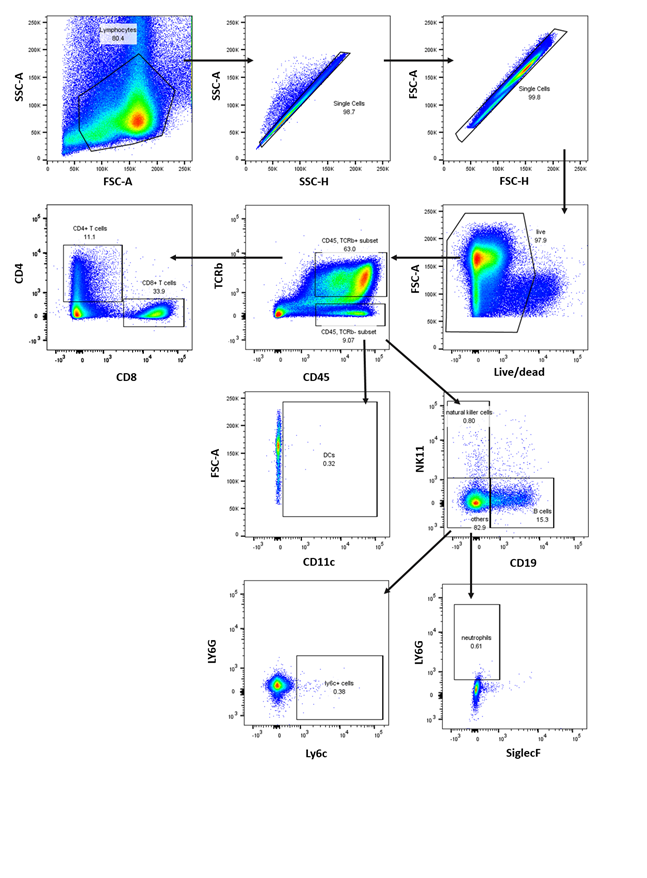
**Fig. S7. Gating strategy for flow cytometric analysis of immune cells in the inguinal lymph node.**

Pseudocolor plots show gating of cells in lymph nodes based on surface markers (CD45, TCRb, CD8, CD4, CD19, CD11c, Ly6C, Ly6G, NK1.1 and siglecF). Percentages indicate frequencies of cells of the parent population.

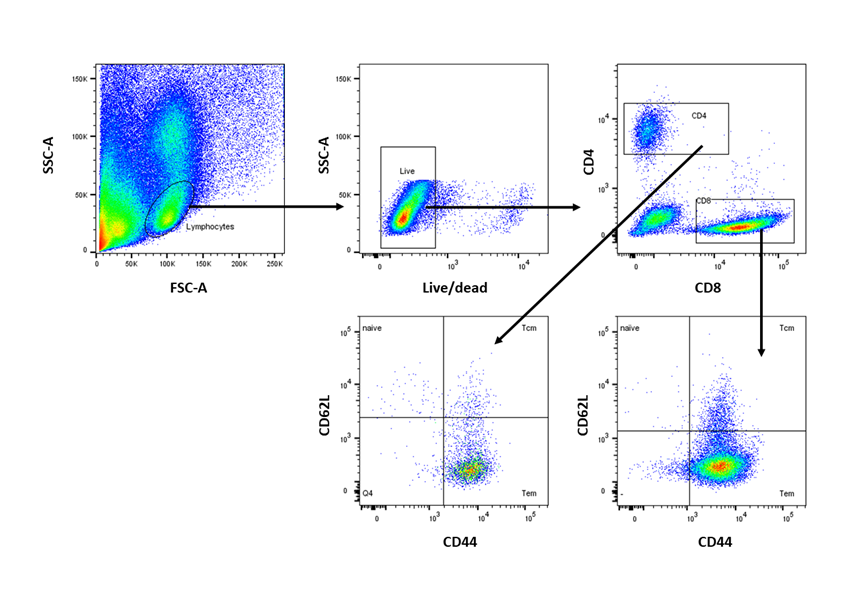

**Fig. S8. Gating strategy for the flow cytometric analysis of T cells in the brain.**

Pseudocolor plots show gating of brain cells based on surface markers (CD4, CD8, CD62L, CD44). T_CM_: central memory T cells; T_EM_: effector memory T cells.

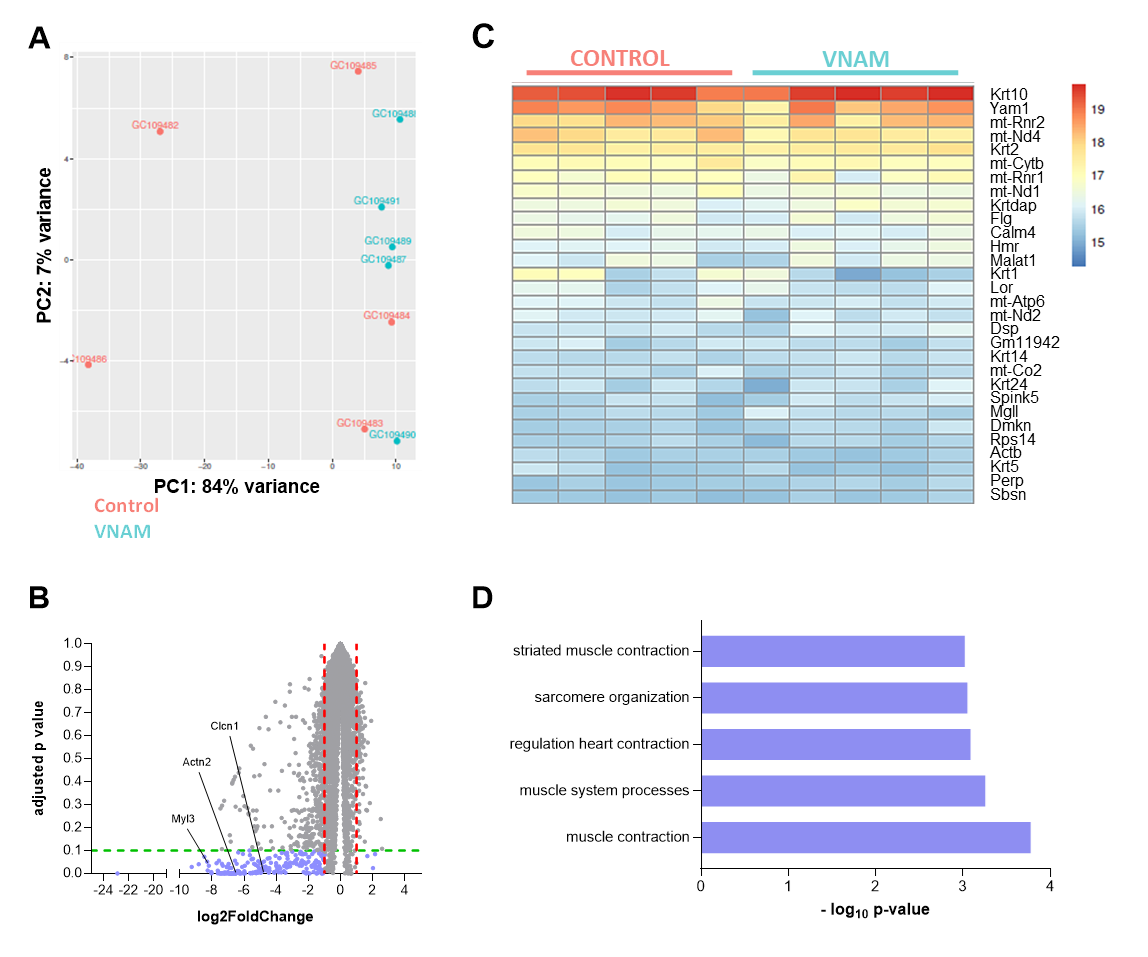
**Fig. S9. Topical VNAM treatment resulted in downregulation of genes involved in muscle contraction processes.**

Mice were topically treated with VNAM or vehicle twice daily for 7 days (n=5 per group). Skin tissue from the footpad was collected after treatment to perform RNA sequencing. (A) PCA plot visualizing sample clustering of VNAM (in blue) and control (in red) samples. (B) Volcano plot showing differentially expressed genes. Fold expression and p values are plotted, each dot represents an individual gene. Genes that were differentially expressed by at least a two-fold difference (red dotted threshold line) and an FDR-corrected p value of <0.1 (green dotted threshold line) are labeled in blue. (C) Heatmap showing the 20 most highly expressed genes in both the VNAM and control group. Data are normalized by variance stabilization formation. (D) Gene ontology and ingenuity pathway analysis tools were used to identify the involved biological processes and pathways upon VNAM treatment**.** FDR; false discovery rate.

**Table S1. Downregulated genes associated with muscle-related processes.**

| **Gene** | **Log2FoldChange** |
| --- | --- |
| Actc1 | -3,5 |
| Actn2 | -6,53 |
| Actn3 | -6,50 |
| Atp2a1 | -6,91 |
| Cacna1s | -7,63 |
| Casq1 | -5,98 |
| Chrnb1 | -2,23 |
| Clcn1 | -4,80 |
| Csrp3 | -7,22 |
| Jsrp1 | -7,01 |
| Klhl41 | -6,28 |
| Ldb3 | -4,46 |
| Lmod2 | -5,11 |
| Lmod3 | -5,26 |
| Mef2c | -2,39 |
| Mybpc1 | -5,25 |
| Mybpc2 | -7,00 |
| Myh1 | -6,53 |
| Myh2 | -7,43 |
| Myh4 | -6,47 |
| Myh7 | -8,19 |
| Myh8 | -4,04 |
| Myl1 | -6,52 |
| Myl2 | -8,27 |
| Myl3 | -8,29 |
| Myl6b | -4,75 |
| Myog | -3,65 |
| Myom1 | -3,84 |
| Mypn | -5,00 |
| Neb | -6,83 |
| Nos1 | -5,53 |
| Pgam2 | -7,28 |
| Ryr1 | -5,59 |
| Six4 | -5,12 |
| Synpo2l | -6,33 |
| Tcap | -6,91 |
| Tmod1 | -3,32 |
| Tnnc1 | -7,38 |
| Tnnc2 | -7,08 |
| Tnni1 | -5,15 |
| Tnni2 | -6,27 |
| Tnnt1 | -6,19 |
| Tnnt3 | -6,81 |
| Tpm1 | -2,8 |
| Tpm2 | -2,17 |
| Trim63 | -4,75 |
| Ttn | -6,69 |
